# Efficacy of Purified Borrelial Lipoproteins (PBL) as an oral formulation in reducing transmission of Lyme spirochetes from reservoir hosts to *Ixodes scapularis* ticks

**DOI:** 10.64898/2026.04.15.718640

**Authors:** Venkatesh Kumaresan, Jolie F. Starling-Lin, Taylor MacMackin Ingle, Nathan Kilgore, J. Seshu

## Abstract

Blocking transmission of *Borrelia burgdorferi* (*Bb*), the causative agent of Lyme disease (LD), from reservoir hosts to humans via *Ixodes scapularis* ticks represents an alternative strategy to reduce LD incidence. Here, we evaluated <u>P</u>urified <u>B</u>orrelial <u>L</u>ipoproteins (PBL) with a combination of adjuvants, for their ability to limit *Bb* transmission using C3H/HeN mice and *Peromyscus leucopus* reservoir models. Immunization with PBL as oral gavage, either alone or nanoparticle-encapsulated, elicited increased antibody responses and reduced pathogen burden in fed larvae and select host tissues. A formulation combining PBL with a recombinant fusion protein adjuvant consisting of <u>C</u>holera Toxin B subunit, <u>O</u>uter surface protein A, and two-tandem repeats of an <u>M</u>-cell-targeting peptide (rCOM) induced durable protective immunity for up to 10 months in C3H/HeN mice. This oral regimen significantly reduced *Bb* burden in host tissues, in fed larvae from vaccinated hosts, molted nymphs, and nymph-challenged naïve mice. Immunization with PBL+rCOM elevated peripheral levels of *Bb*-specific IgG isotypes and increased antigen-specific T cell responses producing IFN-γ and IL-4 at days 28 and 65 post-immunization. Significant protective responses were observed in *P. leucopus*, including strong antibody responses, reduced *Bb* burden in tissues and reduced *Bb* transmission to naïve larvae, independent of sex but influenced by challenge dose. Sodium chloride content in oral formulation modulated vaccine induced protective responses. Notably, *Bb* burden in infected nymphs was reduced during the bloodmeal on vaccinated hosts with decreased pathogen transmission to both vertebrate hosts. These findings support PBL+rCOM as a promising oral, reservoir-targeted, transmission-blocking biologic for controlling Lyme disease.

**Lay Abstract:** Numerous vertebrate hosts serve as reservoirs of pathogens that are transmitted to humans via the bite of blood feeding vectors such as ticks. Lyme disease, caused by *Borrelia burgdorferi* (*Bb*), is the most common tick-borne disease in the US. *Bb* is transmitted to humans following the bite of infected *Ixodes scapularis* ticks. In nature, ticks acquire *Bb* and other pathogens from a variety of reservoir hosts, notably *Peromyscus leucopus*. Therefore, strategies that limit pathogen burden in reservoir hosts or block their transmission via ticks are options to prevent human infectious diseases, circumventing need for human vaccines and therapeutics. An oral, reservoir host-targeted, pathogen-derived, biologic prepared by extracting immunogenic lipoproteins (Purified Borrelial Lipoproteins) from *Bb* and combining them with a mucosal adjuvant derived by fusing Cholera-Toxin B subunit, Outer surface protein A of *Bb* and 2 repeats of an M-cell targeting peptide was tested in C3H/HeN mice and *Peromyscus leucopus* hosts. Single or two dose regimens via the oral route resulted in significant increases in peripheral *Bb* specific antibody responses, select T cell responses, blocking the transmission of *Bb* to naïve *Is* larvae, reducing pathogen burden in vaccinated hosts, and interfering with the infectious cycle of the agent of Lyme disease.

## Introduction

Lyme disease (LD) is the most common tick-borne disease in the United States, with about 86,000 confirmed cases and around 500,000 cases estimated to occur each year, representing a significant public health burden in endemic regions (1, 2). The causative agent, *Borrelia burgdorferi* (*Bb*), is transmitted to humans through the bite of infected *Ixodes scapularis* (*Is*) ticks (3–5). If left untreated, *Bb* can disseminate to deeper tissues and cause severe manifestations, including Lyme arthritis, carditis, and neuroborreliosis (4, 6). Although doxycycline is effective in preventing most severe outcomes, a subset of patients develop Post-Treatment Lyme Disease Syndrome (PTLDS), characterized by persistent symptoms despite antimicrobial therapy (7). The underlying mechanisms of PTLDS remain unclear but may involve persistence of bacterial components such as peptidoglycan, remnants of non-viable spirochetes, coinfections, or host immune dysregulation (8–11). Currently, there is no approved human vaccine, though candidates such as VLA15 (LB6V, VALOR) are in clinical development, and there are a few approved for veterinary use, highlighting the need for alternative strategies to reduce disease incidence (12–14).

Transmission-blocking strategies targeting the natural enzootic cycle of *Bb* offer a promising alternative to human-directed interventions (15–19). The life cycle of *Bb* involves a complex interaction between the tick vector and vertebrate reservoir hosts. Importantly, since there is no transovarial transmission, naïve *Is* larvae acquire *Bb* during their first blood meal from infected reservoir hosts (20). These infected larvae subsequently molt into nymphs, which are primarily responsible for transmitting *Bb* to humans and other naïve reservoir hosts (21). Therefore, reducing pathogen burden in reservoir hosts, the major source of infection for naïve larvae, represents a critical intervention point to disrupt transmission and limit pathogen persistence in nature.

The primary reservoir host for *Bb* in the United States is *Peromyscus leucopus* (white-footed mouse), which harbors multiple tick-borne human pathogens and is capable of infecting 80-90% of ticks feeding on them and plays a central role in maintaining enzootic transmission (22). Notably, *P. leucopus* exhibits immunological features distinct from laboratory *Mus musculus* models, including tolerance to persistent infection (23, 24). This highlights the importance of evaluating interventions in both laboratory and natural reservoir hosts. Targeting reservoir hosts to control these pathogens offers several advantages, including reducing pathogen prevalence at the source and circumventing limitations associated with human vaccines or therapeutics, such as compliance, cost, and potential side effects. While various control strategies including tick-targeted acaricides (25), host population management (26), and reservoir-targeted vaccination (22, 27, 28) have been explored, their effectiveness has been variable due to factors such as early infection in hosts and high background pathogen prevalence.

Previous work has identified several *Bb* antigens capable of inducing protective immune responses targeting multiple hosts- and tick-phase of pathogen. Outer surface protein A (OspA) has been extensively studied and shown to prevent *Bb* transmission by inducing antibodies that neutralize the pathogen within the tick midgut (29–31). Although an OspA-based human vaccine demonstrated efficacy, it was discontinued, and next-generation multivalent formulations are currently under clinical evaluation (12). Outer surface protein C (OspC) can prevent early dissemination in the host (32, 33); however, its high antigenic diversity limits broad protective efficacy (33). A combination of epitopes from multiple OspC proteins in tandem with OspA as a multivalent vaccine has been recently licensed for use in dogs (VANGUARD^®^crLyme) (34, 35). Deploying reservoir-targeted oral vaccines has successfully controlled transmission of several zoonotic diseases (36–39). Reservoir-targeted oral vaccines using OspA delivered via recombinant *Escherichia coli* or vaccinia virus have demonstrated success in reducing *Bb* burden in both laboratory and reservoir hosts, as well as in feeding ticks (18, 40, 41). Notably, long-term field studies using *E. coli*-based vaccines have shown reductions of *Bb* in nymphal infection by about 76%, validating the feasibility of reservoir-targeted approaches (22, 27). A rabies virus vectored Lyme disease vaccines incorporating BB139 (BNSP333-RVG) or OspA (BNSP333-HVG) as inactivated virions were shown to induce robust type1 biased antibody responses (IgG2a/b) and provide short and long-term protection against *Bb* in C3H/HeN mice following intramuscular injections and primarily targeted for development as a human vaccine (42, 43). In addition, *E. coli* expressing OspA, originally formulated as reservoir targeted oral vaccine by Gomes-Solecki et.al (15, 17, 27) was chemically inactivated and formulated as an enteric encapsulated bacterin in pellets was shown to reduce *Bb* infected nymphal ticks in field trials (44). Limitations of current Lyme disease vaccine strategies include (1) reliance on single antigens; (2) requirement for multiple dosing and limited durability of immunity; (3) strain-specific antigenic variation; (4) stage-specific expression of target antigens restricting efficacy across the pathogen life cycle; and (5) perceived safety concerns on live delivery systems and antigen-induced side effects highlighting the need for improved formulations targeting reservoir hosts with a broadened antigenic profile and devoid of live delivery vectors.

An ideal reservoir-targeted vaccine would incorporate multiple antigens expressed during both tick (OspA) and mammalian phases (OspC) of *Bb* infection, thereby inducing broader and more effective protection. In this context, borrelial lipoproteins represent a promising target, as they constitute a major class of immunogenic surface-expressed proteins maintaining the natural antigenic composition. We recently demonstrated that purified borrelial lipoproteins (PBL), derived from *Bb* mutant strains that hyper-express lipoproteins, confer protection when administered intradermally against both needle and tick-mediated challenge and reduce larval acquisition of *Bb* (45). These findings suggest that PBL can induce protective immune responses in reservoir hosts targeting multiple stages of *Bb* enzootic life cycle.

Here, we extend prior formulations based on OspA alone by developing an oral reservoir-targeted vaccine based on array of PBL while leveraging the efficacy of OspA/OspC, among other lipoproteins, in conjunction with mucosal adjuvants (46). Oral vaccination offers practical advantages for large-scale deployment through bait formulations but requires effective strategies to overcome challenges associated with antigen stability and mucosal immune activation, activating both B and T cells for long-term protective immune response. To address this, we evaluated a series of established mucosal adjuvants and delivery systems combined with extraction and purification methods designed to enhance antigen uptake and immune responses via the oral route (47–58). These included oral adjuvants such as Cholera Toxin B subunit (CTB) to activate Th1 responses (59–62), and ligands targeting microfold (M) cells within Peyer’s patches of gut-associated lymphoid tissues (GALT) (63) and delivery systems including polymer-based nanoencapsulation incorporating chitosan and poly(lactic-co-glycolic acid) (64) and ragweed pollen grains (65), which protect antigens from gastric enzymes when delivered in the gastrointestinal tract. Factors such as adjuvant components, targeting strategies, and buffer composition were systematically optimized to improve oral immunogenicity. Using both C3H/HeN mice and *P. leucopus* reservoir host models, we assessed the ability of these formulations to induce protective immune responses, reduce spirochete dissemination, and most critically, to reduce *Bb* transmission to naïve *Ixodes* larvae (66, 67). Our objective was to develop a pathogen-derived, multi-antigen oral vaccine capable of disrupting the tick-pathogen-reservoir axis of Lyme disease. By focusing on reducing pathogen acquisition and transmission at the reservoir-host level, this strategy aims to interfere with the natural enzootic cycle of *Bb* and ultimately reduce human exposure to infected ticks and advances a PBL-based biologic platform to target other tick-borne pathogens.

## Results

### Preparation of Purified Borrelial Lipoproteins (PBL) from *B. burgdorferi*

A key component of this study was to establish a relatively simple and efficient method to extract lipoproteins from *Bb* and evaluate their efficacy to induce protective immune responses as an oral formulation to block transmission of *Bb* between mammalian hosts and naïve *I. scapularis* larvae using murine (C3H/HeN mice) and reservoir host (*Peromyscus leucopus*) models of Lyme disease. As reported previously (68), we extracted borrelial lipoproteins using a detergent (Triton X-114) extraction method separating various borrelial components into the insoluble pellet, aqueous phase (water-soluble proteins), and detergent phase (lipoproteins) proteins. Total stain of proteins separated using an SDS-PAGE gel stained with Coomassie brilliant blue revealed distinct protein profiles between the three phases (Fig 1A). Immunoblot analysis with infection-derived serum, employing a high serum dilution of 1:10,000, revealed the presence of several antigenic proteins present in the detergent phase fraction and in the insoluble pellet compared to the aqueous phase fractions (Fig 1B). Exclusion of non-lipoproteins in TX-114 phase was evaluated by immunoblot analysis using anti-FlaB antibodies. As expected, FlaB (major flagellin of *Bb*) was predominantly detected in the insoluble pellet and aqueous phase fractions, while a faint FlaB band was detected in the detergent phase fraction (Fig 1C). Outer surface Protein C (OspC), a major borrelial lipoprotein contributing to the pathogenic mechanisms of *Bb,* was predominantly detected in the detergent phase (Fig 1D). OspA, OspB, OspC, OspD, and LP6.6 are the top five abundant lipoproteins present in PBL, identified by unbiased proteomics in our previous study (45). The extraction method was used to generate PBL and tested for their efficacy to protect against *Bb* via the intradermal or oral route in conjunction with a variety of immunostimulatory agents in this study.

**Figure 1:**
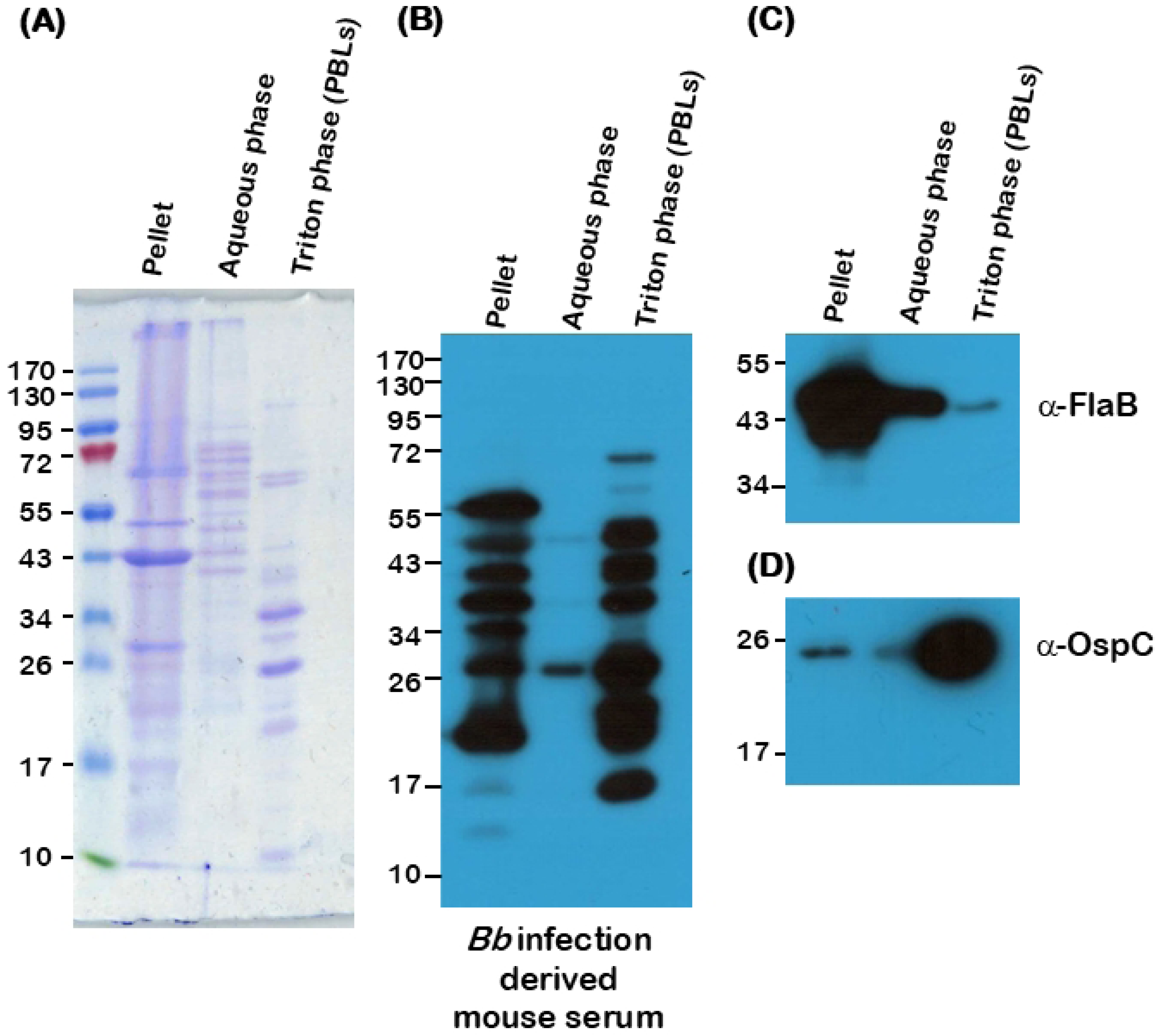
Purified borrelial lipoproteins from *B. burgdorferi*. Total stain of proteins from *Bb* strain B31-A3 partitioned into insoluble pellet (Lane 1), aqueous phase (Lane 2), and detergent phase (Lane 3) using Triton X-114 as the detergent, separated on a 12% SDS-PAGE gel and stained with Coomassie brilliant blue (A). Immunoblots were probed with *B. burgdorferi* infection-derived serum (B), anti-FlaB (C) and anti-OspC (D). Molecular masses of protein standards in kilodaltons are indicated to the right.

### Oral administration of PBL induces protective immune responses in C3H/HeN mice

Purified Borrelial Lipoproteins (PBL) and purified recombinant Outer surface protein A (rOspA) were used to immunize C3H/HeN mice (*n*=3) orally on day 0 and day 30, with a dosage of 100 μg/mouse (Fig 2A), either alone or in combination with immunomodulatory agents as detailed in Fig 2B. Oral immunization with two doses of PBL without any adjuvant induced significantly higher antibody responses at day 42 post-immunization compared to naïve controls when *Bb* lysates or PBL were used as coating antigens while at day 28 there was no significant increase in titers following oral administration (Fig 2C-E). As anticipated, levels of antibodies were higher in mice immunized via the intradermal route, reacting with additional antigens (Fig 2C-F). Moreover, PBL administered orally following encapsulation as chitosan-PEG-PLGA nanoparticles induced antibody responses in two out of three mice at day 42, with a similar antibody response following oral administration of PBL (Fig 2E-F). Immunoblot analysis of serum from mice vaccinated with different preparations and routes revealed that mice with lower end-point titers exhibited reduced reactivity to total *Bb* lysates derived from spirochetes propagated under conditions mimicking either unfed (23 °C, pH 7.6) or fed (37 °C, pH 6.8) tick midgut environments. (Fig 2F). No viable spirochetes were detected in different tissues in mice that exhibited seroconversion following experimental infection via needle challenge with 1x10^5^ *Bb* strain B31-A3/mouse and was found to correlate with levels of antibodies induced (Fig 2G). One out of four mice vaccinated with PBL as nanoparticles with lower seroconversion was positive for *Bb* in several tissues (Fig 2G). Viable spirochetes were detected in all tissues from unvaccinated mice, as expected, serving as positive controls for infection. qPCR analysis showed higher levels of *Bb*-specific DNA in the skin of unvaccinated controls and in mice without seroconversion, whereas lower levels were detected in the skin and joints of seroconverted mice, likely reflecting both viable and non-viable spirochetes (Fig 2H, I). Notably, PBL-ID vaccinated mice showed no detectable *Bb*-specific DNA in skin and joints (Fig 2H, I), suggesting that the superior protective immune response of PBL is administration via the intradermal route. These studies provide a proof of principle of the vaccination strategy to use PBL as oral preparations and sets the stage for evaluating a series of formulations to increase the efficacy of protection via oral route.

**Figure 2:**
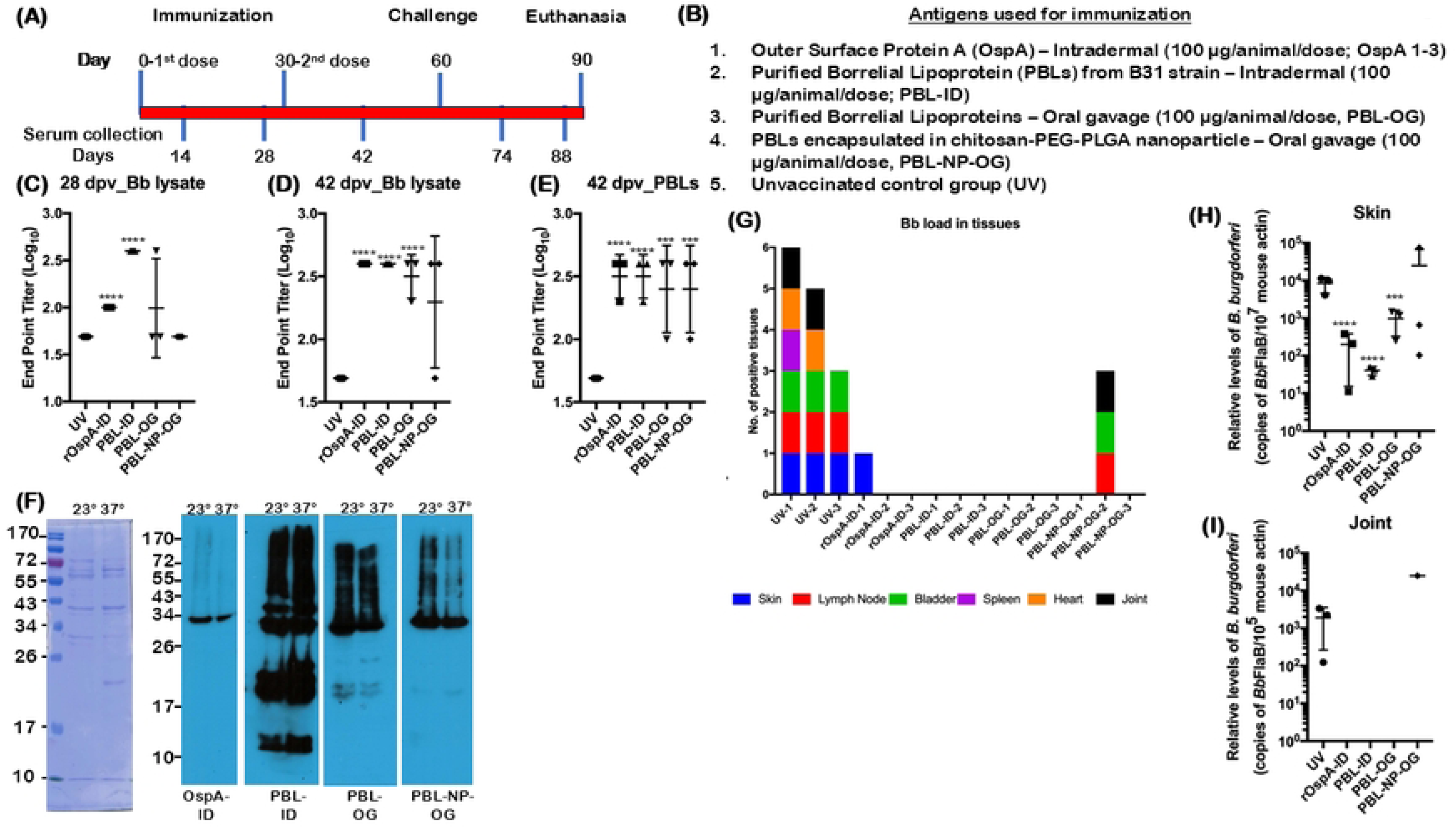
Efficacy of protection conferred by PBL in C3H/HeN mice against challenge with *B. burgdorferi* strain B31-A3. Schematic representation showing the timeline for immunization, challenge, euthanasia and serum collection time points (A) and formulations, routes and dosage used for immunization (B). Endpoint titers of serum antibodies from immunized mice at day 28 (C) and day 42 post-vaccination against total *Bb* lysate (D) or PBL (E), determined by ELISA. Immunoblot analysis of serum from mice at day 28 post-immunization (F). Stacked column graph showing viable spirochetes propagated in BSKII medium from tissues isolated from challenged mice at 28 day post-infection (G). Quantitative real time PCR analysis of spirochetal burden in skin (H) and joint (I) tissues following challenge. Copies of borrelial *FlaB* normalized to mouse b-actin copies. Comparisons between vaccinated groups and unvaccinated controls were performed using a two-tailed Student’s *t*-test. Statistical significance is indicated as: *p < 0.05, **p < 0.01, ***p < 0.001, ****p < 0.0001.

### Oral vaccination with PBL reduced the transmission of *Bb* from mice to naïve ticks

A second study with PBL formulated with immunomodulatory agents to enhance the levels of protective capability as an oral formulation was compared with PBL alone, administered orally or intradermally (Fig S1A, B). The ability of naïve larvae to acquire *Bb* was determined from mice challenged via needle following an immunization schedule, number of doses, and concentration of PBL per dose used in the previous experiment (Fig S1A B). At 55 days post-vaccination, all mice were needle-challenged with 1x10^5^ spirochetes (*Bb* strain B31-A3) per mouse, and tick feeding was initiated at day 88 (33 dpi) with euthanasia at day 118 (Fig S1A). Mice vaccinated twice with PBL via intradermal or via an oral gavage exhibited significantly higher seroconversion compared to unvaccinated controls at day 14, 28 and 42 (Fig S1C-F) and found to have kinetics as observed in the first experiment (Fig 2). Importantly, naïve larvae fed on PBL-vaccinated mice had significantly lower *Bb* burden compared to those from unvaccinated mice (Fig S1G). Mice vaccinated with rOspA intradermally served as a control for levels of transmission of *Bb* to naïve *Is* larvae which was significantly lower, as expected, compared to unvaccinated controls (Fig S1G). qPCR analysis revealed that *Bb* burden in skin of PBL-OG mice was not significantly lower (Fig S1H); however, the levels in joints showed a significant reduction in mice vaccinated with PBL orally or as nanoparticles (Fig S1I). Viable spirochetes were detected from several tissues when propagated in BSKII media following euthanasia in unvaccinated and several of the vaccinated mice (Fig S1J). One key difference between experimental results reported in Fig 2 and Fig S1 is the recovery of viable spirochetes in select tissues which are likely contributed by the difference in the duration of infection which was for 30 days after challenge in first experiment (Fig 2) while 63 days in the second experiment (Fig S1). Collectively, both independent experiments demonstrated the critical immune correlates of protection and the ability to block pathogen transmission to naïve larvae from immunized mice via oral PBL-based formulations. Although oral formulations via PBL with or without nanoparticles induced significant protective immune response, nanoparticle-based delivery did not significantly elevate the immune responses, suggesting the need for additional oral adjuvants.

### Optimization of PBL-based oral vaccination platform for blocking transmission of *Bb* to naïve *Is* larvae

To enhance the oral immunogenicity of PBL, we developed a series of recombinant proteins incorporating components designed to modulate mucosal immune responses. First, we targeted microfold (M) cells in Peyer’s patches, which exhibit high transcytosis activity within the intestinal mucosa, making them attractive targets for M-cell-binding ligands to enhance antigen uptake and promote antigen-specific immune responses (69). Since OspA is a major vaccine candidate, we constructed a plasmid using the pET23a backbone to express a recombinant fusion protein rOspA-C2X-M-cell (Fig S2A). This construct included a 6× histidine tag at the N-terminus and encoded *Bb* outer surface protein A (rOspA) fused to two tandem repeats of an M-cell binding peptide at the C-terminus. Flexible linker sequences (GGGGS) were inserted between the peptide repeats to promote proper folding of the fusion protein, thereby enhancing its stability and immunogenicity for oral vaccination (Fig S2A). A ∼31.2 kDa rOspA fused to M-cell binding peptide was purified and used for immunization (Fig S2A). This plasmid construct was further modified by cloning the sequence for a mucosal adjuvant, Cholera Toxin B subunit (CTB), at the N-terminus to generate a construct resulting in a fusion protein rCTB-OspA-C2X-M-cell (Fig S2B). The immunomodulatory products produced from these two plasmids are referred to as rOM (<u>rO</u>spA-C2X<u>-M</u>-cell-binding peptide) and rCOM (rCTB-rOspA-C2X-M-cell binding peptide) for the remainder of this manuscript.

### Efficacy of protection conferred with <u>r</u>ecombinant <u>O</u>spA-<u>M</u>-cell binding peptide (rOM)

To evaluate the efficacy of rOM as a standalone vaccine, we immunized 4-week-old C3H/HeN mice with three oral doses of rOM alone or formulated with either the mucosal adjuvant CTB or the synthetic polymer Eudragit, which enhances antigen retention in the gastrointestinal tract (64). Vaccinated mice were subsequently challenged with infectious *B. burgdorferi* B31-A3. (Fig S3A and B). While rOM with rCTB or Eudragit exhibited significant levels of seroconversion in all the vaccinated mice at day 35, other immunogens such as rOspA or rOM alone administered orally did not (Fig S3C and D). Despite seroconversion, most of the vaccinated mice transmitted *Bb* to the tick larvae (Fig S3E). All mice except one, vaccinated with rOM+CTB+Eudragit, were positive for *Bb* in the tissues tested for the presence of spirochetes in BSKII medium (Fig S3F) or by qPCR of the skin and joint (Fig S3G, H). Overall, these observations suggested that oral formulations with rOM and CTB as adjuvant elicited higher antibody response against OspA in the *Bb* lysate in mice, although the overall immune response was not sufficient to confer protection against *Bb* infection or reduce the transmission levels to naïve *Is* larvae. These findings suggest that incorporating CTB with rOM as an adjuvant in PBL oral formulations may enhance protective immune responses against *Bb* in reservoir hosts and reduce transmission to naïve *Ixodes scapularis* larvae.

### Efficacy of protection conferred with <u>r</u>ecombinant CTB-OspA-M-cell binding peptide (rCOM)

Building on prior findings demonstrating the ability of CTB to induce robust antibody responses, we generated a fusion protein in which recombinant CTB was linked to OspA and an M-cell–binding peptide (rCOM; Fig 2C-E). We then immunized six groups of four-week-old C3H/HeN mice using different formulations. Mice vaccinated intradermally with rOspA served as a positive control (Group 1), followed by intradermal PBL (PBL-ID; Group 2) and oral gavage PBL (PBL-OG; Group 3). To enhance oral delivery, PBL were also administered using ragweed pollen grains as a delivery vehicle (PBL-RPG-OG; Group 4). To evaluate rCOM as an oral adjuvant, mice were immunized with PBL+rCOM (PBL+rCOM-OG; Group 5), and to assess its standalone efficacy, rCOM alone was administered orally (rCOM-OG; Group 6). Significant antibody responses were observed on days 28, 42 (data not shown), and 74 post-vaccination in three out of four mice immunized with PBL-RPG-OG or PBL+rCOM-OG (Fig 3C). Intradermal vaccination with PBL or rOspA induced robust antibody responses in all animals. In contrast, PBL-OG induced seroconversion in only two out of four mice, whereas PBL-RPG-OG and PBL+rCOM-OG achieved seroconversion in three out of four mice. Oral administration of rCOM alone did not induce seroconversion (Fig 3C).

**Figure 3:**
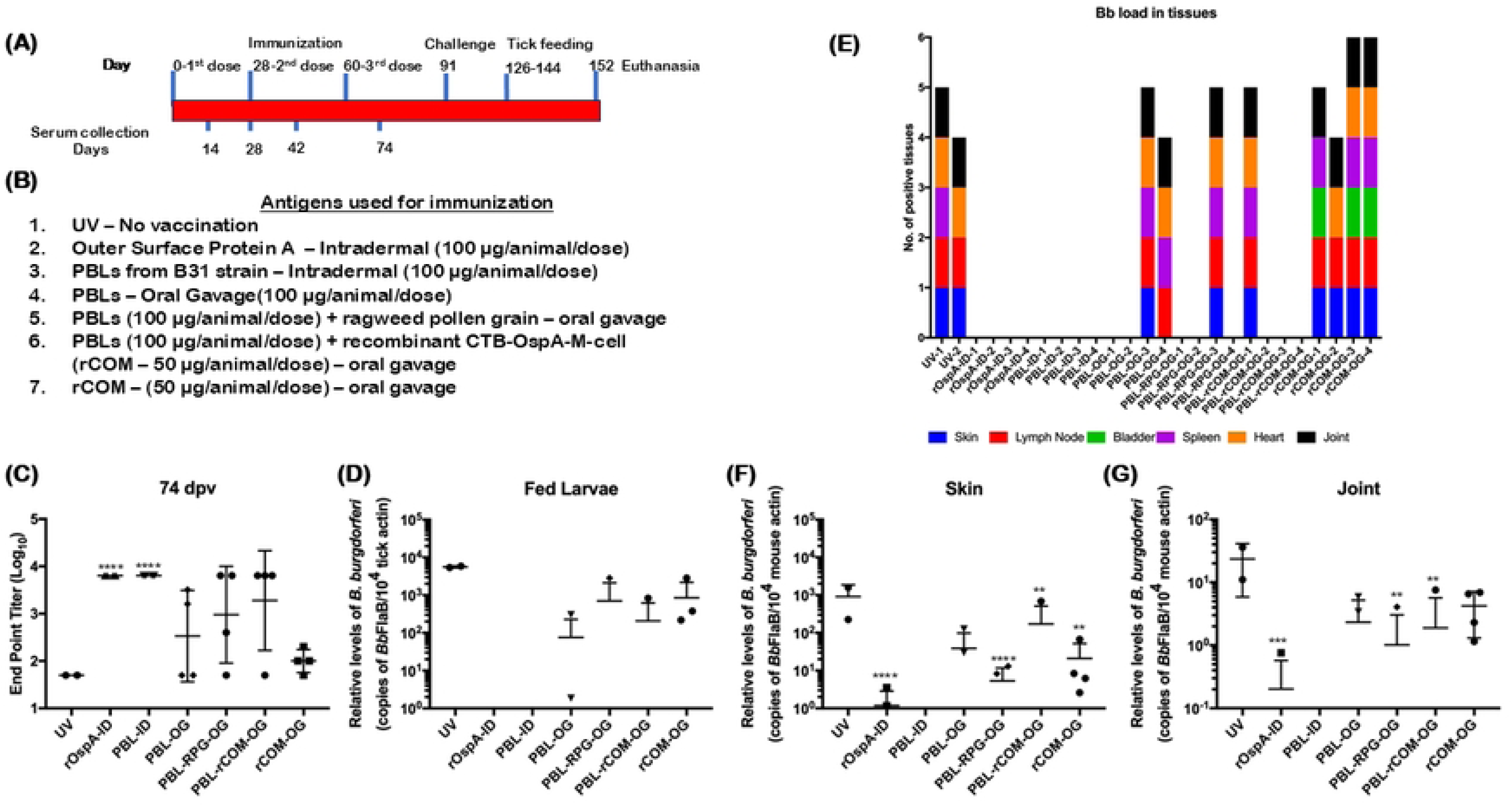
Levels of induction of antibody responses against *Bb* following three oral gavages with PBL plus rCOM. Schematic representation showing the timeline for immunization, challenge, euthanasia and serum collection time points (A). Formulations, routes and dosage used for immunization (B). Endpoint titers of serum antibodies from immunized mice post-immunization against total *Bb* lysate determined by ELISA at day 74 post-vaccination (C). Quantitative real time PCR analysis of spirochetal burden in larvae fed at day 126 with copies of borrelial *FlaB* normalized to tick actin (D). Stacked column graph showing the presence of *Bb* in six tissues (mentioned at the bottom of the graph) of mice on euthanasia at day 152 (E). Quantitative real time PCR analysis of spirochetal burden in skin (F) and joints (G) at day 152 with copies of borrelial *FlaB* normalized to mouse b-actin copies. Comparisons between vaccinated groups and unvaccinated controls were performed using a two-tailed Student’s *t*-test. Statistical significance is indicated as: *p < 0.05, **p < 0.01, ***p < 0.001, ****p < 0.0001.

Consistent with seroconversion, both PBL-RPG-OG and PBL+rCOM-OG reduced *Bb* acquisition by tick larvae, with larvae from three out of four mice showing no detectable *Bb*. Intradermal administration of rOspA or PBL completely blocked transmission compared to unvaccinated controls (Fig 3D). Similarly, viable spirochetes were recovered from all tissues of unvaccinated and rCOM-OG mice following culture in BSKII medium, whereas only one out of four mice in the PBL-RPG-OG and PBL+rCOM-OG groups showed spirochete outgrowth (Fig 3E). qPCR analysis revealed significantly lower levels of *Bb*-specific DNA in the skin and joint tissues of mice vaccinated with PBL-RPG-OG and PBL+rCOM-OG compared to unvaccinated mice or those receiving rCOM-OG, consistent with seroconversion patterns (Fig 3F, G). The single mouse in each of the PBL-RPG-OG and PBL+rCOM-OG groups with detectable *Bb* burden also exhibited spirochete outgrowth in multiple tissues. It is important to note that *Bb* DNA detected in certain tissues, particularly skin, may partially reflect presence of non-viable spirochetes.

Importantly, the low seroconversion, high transmission rates, and widespread recovery of viable spirochetes in the rCOM-OG group indicate that OspA delivered as part of rCOM alone is insufficient to drive protective immunity via oral route. Rather, the combination of PBL and rCOM is required for effective protection. Given the comparable efficacy of PBL+RPG-OG and PBL+rCOM-OG, and the advantages of recombinant protein-based formulations, subsequent studies focused on PBL+rCOM-OG. These findings guided further optimization of this formulation and refinement of the immunization strategy to maximize immune responses and limit *Bb* transmission to naïve ticks.

### Oral PBL+rCOM Vaccination Reduces *B. burgdorferi* Across Tick Developmental Stages

To further validate the protective efficacy of PBL+rCOM without additional immunostimulatory components, we immunized 4-week-old C3H/HeN mice with two doses of PBL+rCOM administered either intradermally or via oral gavage, with rCOM-ID serving as a control (Fig 4A, B). Compared to unvaccinated controls, mice vaccinated with PBL+rCOM by either route exhibited significantly higher antibody titers at days 28 and 42 post-vaccination, while rCOM-ID induced less significant antibody titers (Fig 4C, D). Correspondingly, acquisition of *Bb* by naïve *Ixodes scapularis* larvae feeding on these mice was markedly reduced (Fig 4E), consistent with the absence of viable spirochetes in host tissues (Fig 4F). qPCR analysis showed no detectable *Bb*-specific DNA in the skin of most vaccinated mice, with only two PBL+rCOM-OG vaccinated mice exhibiting low levels, significantly lower than those observed in unvaccinated controls (Fig 4G). Similarly, *Bb* DNA was undetectable in the joint tissues of vaccinated mice, in contrast to unvaccinated controls (Fig 4H), correlating with high seroconversion, less larval acquisition and absence of viable spirochetes in tissues.

**Figure 4:**
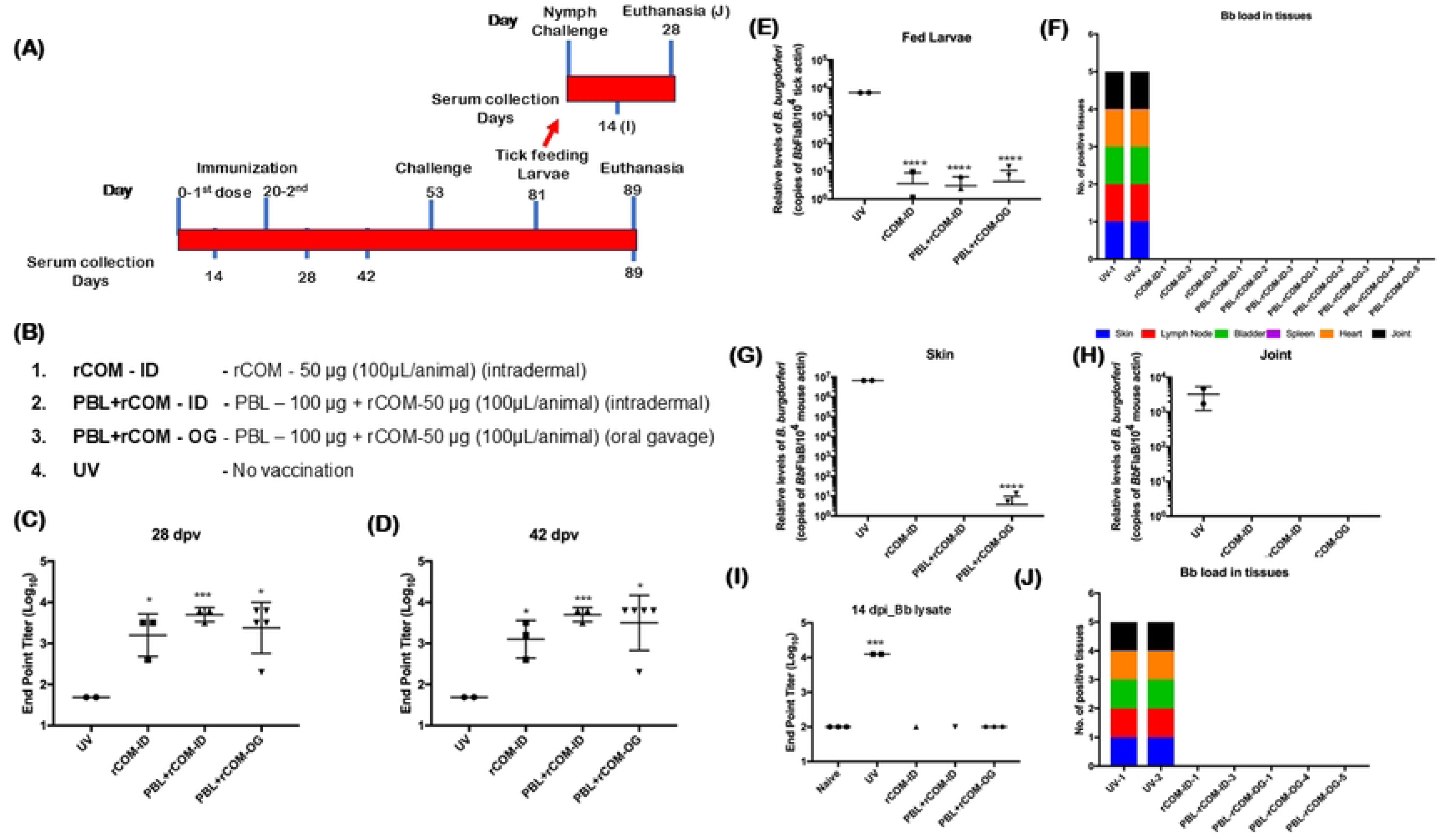
Level of induction of antibody responses against *Bb* following two oral gavages with PBL plus rCOM. Schematic representation showing the timeline for immunization, challenge, euthanasia and serum collection time points (A). Formulations, routes and dosage used for immunization (B). Endpoint titers of serum antibodies from immunized mice post-immunization against total *Bb* lysate determined by ELISA at day 28 (C) and day 42 (D) post-vaccination. Quantitative real time PCR analysis of spirochetal burden in larvae fed at day 81 with copies of borrelial *FlaB* normalized to tick actin (E). Stacked column graph showing the presence of *Bb* in six tissues (mentioned at the bottom of the graph) of mice on euthanasia at day 89 (F). Quantitative real time PCR analysis of spirochetal burden in skin (G) and joints (H) at day 89 with copies of borrelial *FlaB* normalized to mouse b-actin copies. Endpoint titers of serum antibodies from mice challenged with nymphs collected from vaccinated and unvaccinated mice at day 14 post-challenge (I) and stacked column graph showing the presence of *Bb* in six tissues (mentioned at the bottom of the graph) of mice challenged with nymphs on euthanasia at day 28 post-challenge (J). Comparisons between vaccinated groups and unvaccinated controls were performed using a two-tailed Student’s *t*-test. Statistical significance is indicated as: *p < 0.05, **p < 0.01, ***p < 0.001, ****p < 0.0001.

To assess transmission potential, nymphs molted from larvae that had fed on vaccinated or control mice were used to challenge naïve C3H/HeN mice. Three nymphs from PBL+rCOM-OG mice, one each from PBL+rCOM-ID and rCOM-ID groups, and two from unvaccinated mice were used. Serological analysis at 14 days post-infection showed that nymphs derived from vaccinated mice did not induce *Bb*-specific antibody responses in naïve hosts, whereas nymphs from unvaccinated mice induced robust seroconversion (Fig 4I). Consistently, viable *Bb* was recovered only from mice challenged with nymphs originating from unvaccinated controls (Fig 4J). Collectively, these findings demonstrate that immunization with PBL, combined with CTB fused to OspA and an M-cell binding peptide (rCOM), induces strong protective immunity in C3H/HeN mice and effectively reduces *Bb* transmission across multiple stages of the tick enzootic cycle.

### Oral PBL+rCOM vaccination induces peripheral and mucosal antibody responses against borrelial antigens

C3H/HeN mice were orally vaccinated with PBL+rCOM, with a booster dose administered on 14 dpv. To determine the antibody subtypes in the peripheral blood, serum samples were collected from mice on 28 dpv, and anti-mouse antibodies, including total Ig, IgG, IgG1, IgG2a, IgG2b, IgG3, IgM, and IgA, were quantified against borrelial lysate. Results showed that PBL+rCOM vaccinated mice exhibited a significantly high concentration of total Ig and IgG compared to unvaccinated mice, with a less significant IgM response and no difference in IgA observed (Fig 5A). Further IgG subclass analysis revealed significantly elevated IgG1, IgG2a, and IgG2b responses in vaccinated mice, with comparatively smaller differences observed for IgG3 between vaccinated and control groups (Fig 5B). These results indicate that PBL+rCOM elicits a strong systemic humoral immune response against borrelial antigens.

**Figure 5:**
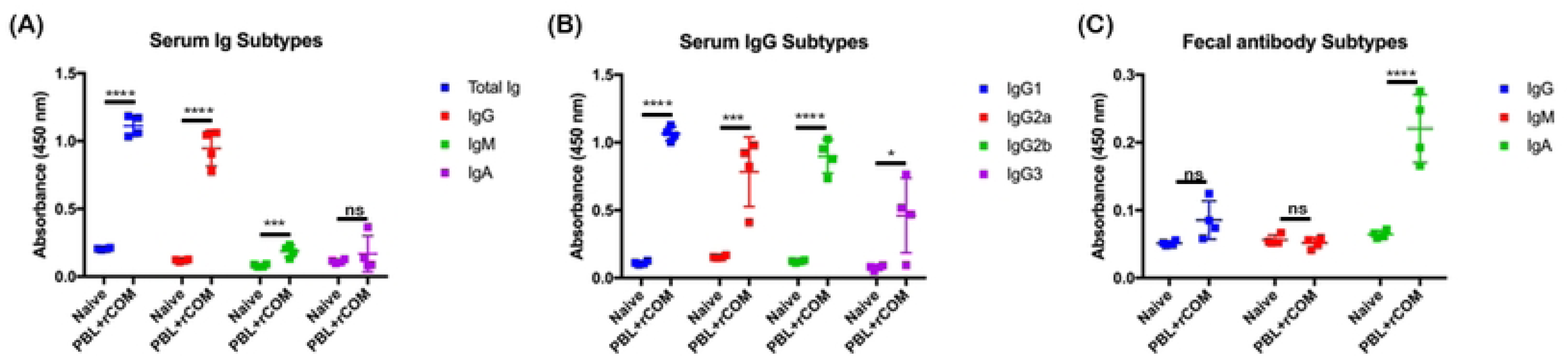
Serum and fecal antibody isotypes following oral gavage with PBL+rCOM. Graphs showing the absorbance (OD450) as a measure of different isotypes of serum antibodies binding to total *Bb* lysate for total Ig, IgG, IgM and IgA antibodies (A) and IgG subtypes IgG1, IgG2a, IgG2B and IgG3 (B) from the mice sera collected at 28 dpv. Graph showing the absorbance (OD450) as a measure of different isotypes (IgG, IgM and IgA) of antibodies binding to total *Bb* lysate from the feces collected at 28 dpv (C) determined by ELISA. Comparisons between vaccinated groups and unvaccinated controls were performed using a two-tailed Student’s *t*-test. Statistical significance is indicated as: *p < 0.05, **p < 0.01, ***p < 0.001, ****p < 0.0001.

Mucosal antibody responses were assessed using wet fecal samples collected at day 28 post-vaccination and analyzed against borrelial lysate. A high concentration of IgA antibodies was detected in feces relative to other isotypes (Fig 5C). In contrast, IgG and IgM levels in fecal samples did not differ significantly between vaccinated and control groups, as expected for mucosal antibody profiles. These findings suggest that oral PBL+rCOM vaccination induces robust secretory IgA responses even at 28 dpv in the gastrointestinal tract, implying a prolonged immune response against *Bb* antigens likely mediated by the mucosal adjuvant effects of the Cholera Toxin B subunit.

### Induction of T cell memory response by oral PBL+rCOM against borrelial lipoproteins

To evaluate the efficacy of PBL+rCOM vaccination in inducing T cell memory responses against borrelial lipoproteins, C3H/HeN mice were immunized via oral gavage with a booster dose administered at day 14 post-vaccination. Splenocytes were collected at days 28 and 65 post-vaccination, and ELISPOT assays were performed to quantify PBL-specific IFN-γ producing Th1 cells and IL-4 producing Th2 cells (Fig 6A). Mice vaccinated with PBL+rCOM exhibited significantly higher frequencies of IFN-γ producing Th1 cells at both day 28 and day 65 post-vaccination (Fig 6B, C), as well as increased IL-4 producing Th2 cells at day 65 (Fig 6D), compared to unvaccinated controls. Results are presented as spot-forming units (Fig 6B-D, bottom panels). These findings indicate that oral vaccination with PBL+rCOM effectively induces borrelial lipoprotein-specific memory T cell responses, which likely contribute long-term protective efficacy with this immunization strategy.

**Figure 6:**
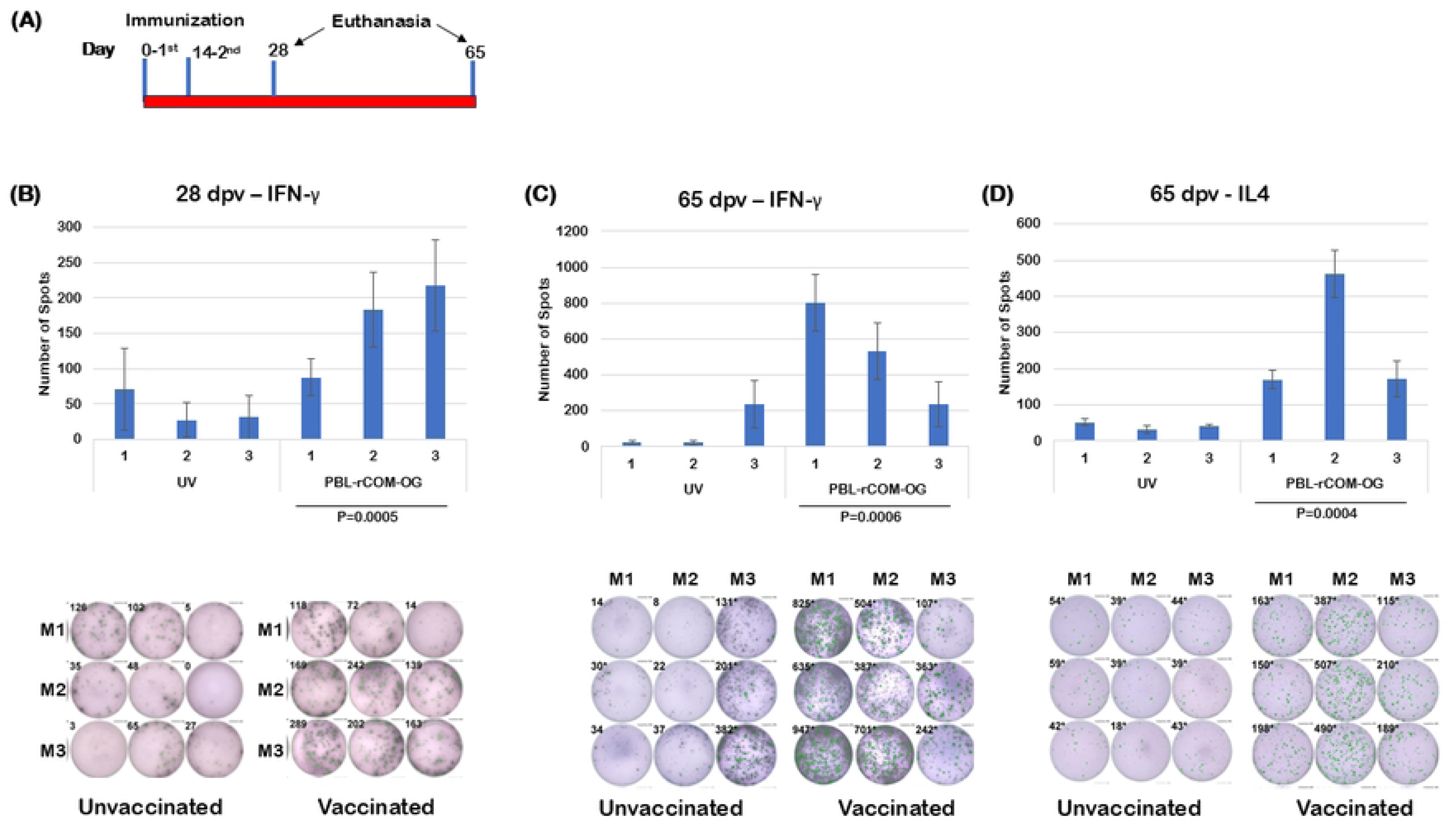
Th1 and Th2 response induced following oral gavage with PBL+rCOM determined by ELISPOT. Schematic representation showing the timeline for immunization, euthanasia and serum collection time points (A). Graphs showing the number of spots of interferon-gamma producing splenocytes at 28 dpv (B) and 65 dpv (C) and interleukin-4 producing splenocytes at 65 dpv (D). The images at the bottom of each graph show the spots in each well directed against PBL. Comparisons between vaccinated groups and unvaccinated controls were performed using a two-tailed Student’s *t*-test. Statistical significance is indicated as p values below the bar graphs.

### Levels and length of protection conferred by a single oral dose of PBL+rCOM

To further advance the utility of PBL+rCOM as a reservoir host targeted oral biologic to interfere with the transmission cycle of *Bb* from vertebrate hosts to naïve ticks, we vaccinated a group of 10 (*n*=10) four-week old C3H/HeN mice (numbered 4 to 13) with a single dose of PBL+rCOM and collected serum at different time points post-vaccination and challenged with *Bb* strain B31-A3 at 1x10^5^ *Bb*/mouse at day 302 (Fig 7A). Three age-matched, sex-matched, C3H/HeN mice (numbered 1 to 3) served as unvaccinated controls. Compared to serum from control mice, all vaccinated mice had higher antibody titers against PBL at day 14, 28, 105 and 223, suggesting that a single oral dose of PBL+rCOM, despite variations in titers between vaccinated mice, was capable of inducing antibody responses over a prolonged period of time (Fig 7B). A correlation was also observed in the levels of immunoreactivity of the serum with total borrelial lysate by immunoblot analysis at day 302, with mice numbers 7, 8 and 9 having lower antibody responses compared to mice numbers 4, 5, 6, 10, 11, 12, and 13 (Fig 7C, D). Notably, levels of acquisition of *Bb* by naïve larvae fed on PBL-rCOM vaccinated mice were significantly (p<0.0001) lower than that from unvaccinated mice (Fig 7E). While vaccinated mice numbers 4, 5, and 11 had higher *Bb* burden in skin, all other mice had lower relative burden of *Bb* (*flaB* copies) compared to unvaccinated controls (Fig 7F). Relative *Bb* burden in the joints of the vaccinated mice was significantly lower than unvaccinated mice (p<0.05 Fig 7G). While mice numbers 11 and 12 had higher antibody levels, the presence of a higher *Bb* burden in skin and joints could be attributed to amplification of *flaB* from dead spirochetes, as the levels of transmission of *Bb* to naïve *Is* larvae from these two mice were lower than unvaccinated mice or those with lower antibody titers (Fig 7B-G). Overall, these studies demonstrate that oral vaccination with a single dose of PBL+rCOM significantly reduces the transmission of *Bb* to naïve ticks, which is a key determinant of the efficacy sought for this formulation with a protective response extending to around 9 months as an oral reservoir-targeted biologic against agent of Lyme disease.

**Figure 7:**
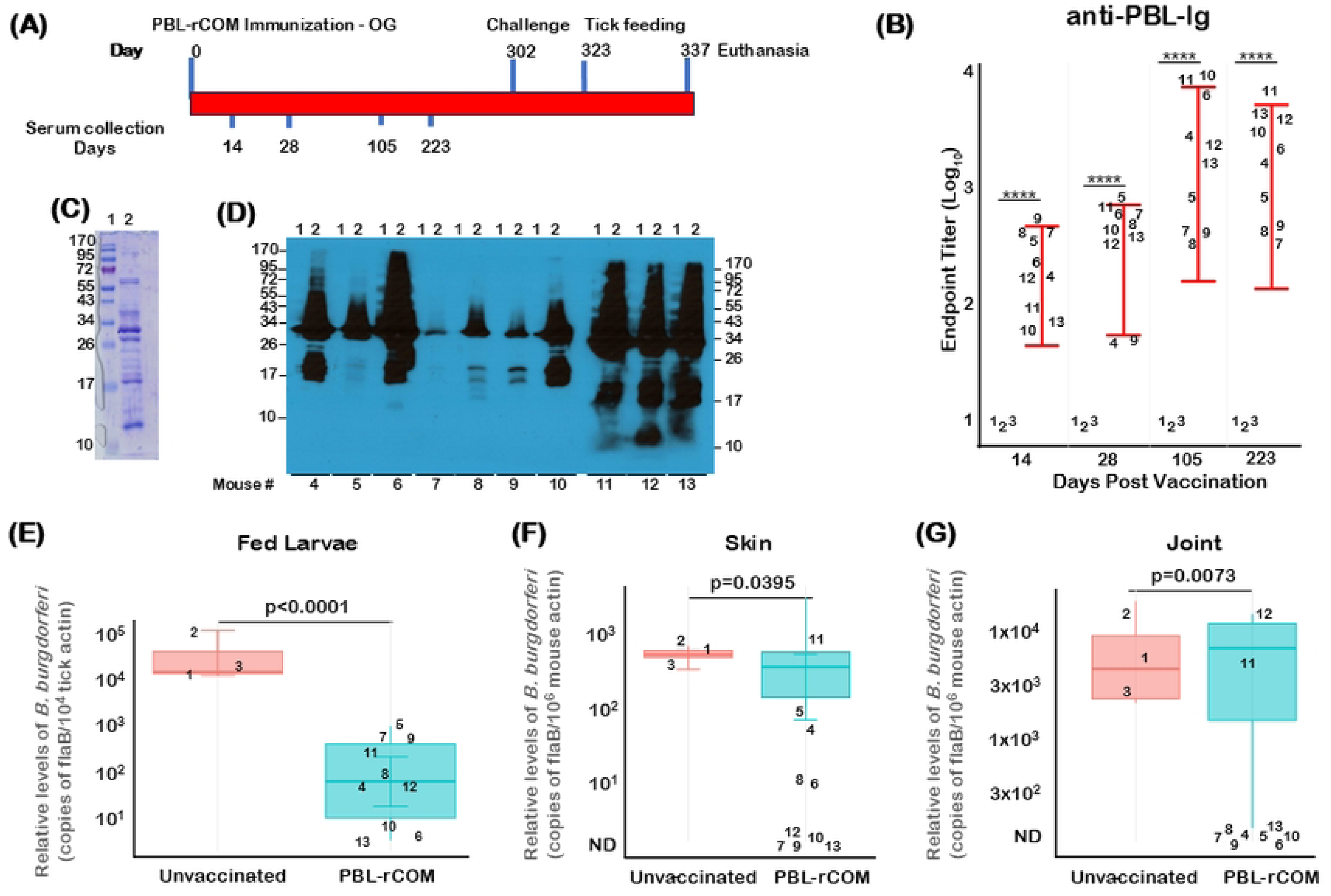
Efficacy of protection conferred by a single oral dose of PBL+rCOM. Schematic representation showing the timeline for immunization, challenge, serum collection and euthanasia time points (A). Endpoint titers of serum antibodies from mice at day 14, 28, 105 and 223 (B). Coomassie blue stained gel of PBL (C) and immunoblot analysis of serum from each of the vaccinated mice at day 223 showing reactivity to PBL (D). Quantitative real time PCR analysis of spirochetal burden in fed larvae - copies of borrelial *flaB* gene normalized to tick actin copies (E) and quantitative real time PCR analysis of spirochetal burden in skin (F) and joint (G). Note significant reduction in *Bb* burden in ticks from mice vaccinated following a single oral administration of the PBL+rCOM. Comparisons between vaccinated groups and unvaccinated controls were performed using a two-tailed Student’s *t*-test. Statistical significance is indicated as p values.

### Effects of oral vaccination with PBL+rCOM on the mouse-tick-mouse cycle of *Bb* infection

To check if the nymphs from long-term vaccinated mice still exhibit reduced transmission, larvae fed on mice vaccinated with PBL+rCOM-OG (Fig 8A, host C) were allowed to molt into infected flat nymphs and allowed to feed on naïve C3H/HeN mice (Fig 8A, host D). Serum was collected after infected nymphs fed to repletion (Fig 8A, host E) and end point titers against PBL were determined. As shown in Figure 8B, there were higher titers in naïve C3H/HeN mice infected with nymphs from unvaccinated M2, vaccinated M4, and M13 compared to naïve mouse. On the other hand, the titers were lower when infected with nymphs from vaccinated M6, demonstrating that there was lower *Bb* burden in these ticks, similar to that in the naïve control.

**Figure 8:**
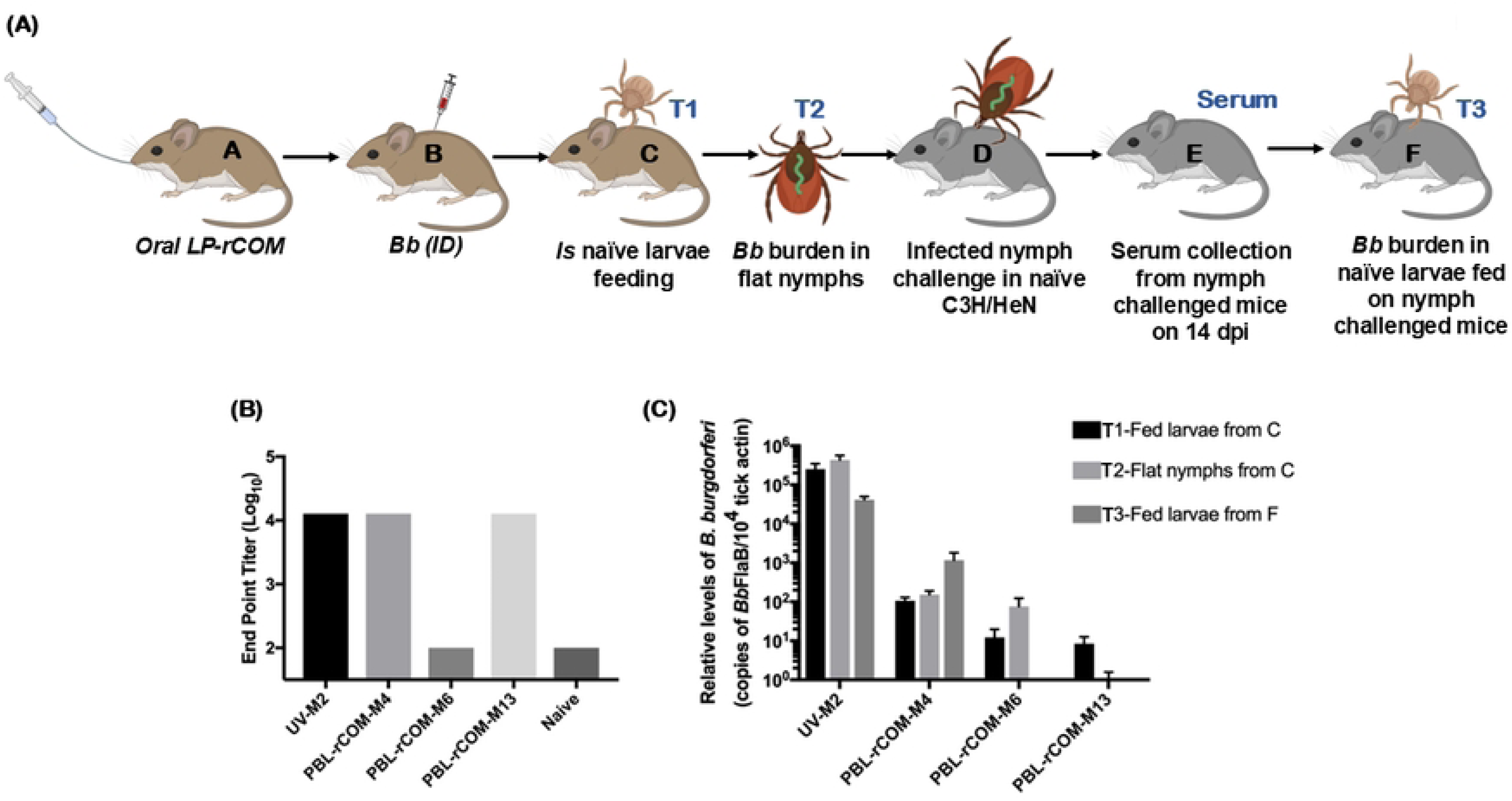
Efficacy of single oral PBL+rCOM dose in reduction in transmissibility of *Bb* via *I. scapularis* ticks. Schematic of mouse-tick-mouse infectious cycle using ticks obtained following needle challenge with vaccinated (M4, M6 and M13) and unvaccinated (M2) mice (A). End point titers of serum from mice challenged with infected flat nymphs from M2, M4, M6 and M13 as well as from naïve control mice (B); Note reduced titers in M6 similar to naïve control indicating lack of colonization of *Bb* reflecting protective efficacy of PBL+rCOM (B). Quantitative real time PCR analysis of spirochetal burden in 1) fed larvae from host C, 2) Flat nymphs from host C, and 3) Fed larvae from host F (C). Copies of borrelial *flaB* gene normalized to tick actin copies. Comparisons between vaccinated groups and unvaccinated controls were performed using a two-tailed Student’s *t*-test. Statistical significance is indicated as: *p < 0.05, **p < 0.01, ***p < 0.001, ****p < 0.0001.

To evaluate the impact of a single oral dose of PBL+rCOM on *Bb* transmission, fed larvae from vaccinated and unvaccinated C3H/HeN mice were tracked through their developmental stages (Fig 8A). qPCR was used to determine the *Bb* burden in ticks at each stage (Ref – Fig 8A: Fed larvae, T1-from mouse C; flat nymphs, T2 from mouse C; and fed larvae, T3 from mouse F) and at least five ticks were used for the quantification. Results showed that fed larvae (T1) from the unvaccinated mouse (UV-M2) harbored high *Bb* burden, which persisted in flat nymphs (T2) and in fed nymphs following feeding on naïve mice, and was transmitted to subsequent naïve larvae (T3) (Fig 8C). In contrast, larvae from vaccinated mice (PBL+rCOM-M4) showed reduced *Bb* burden at all stages, including fed larvae, flat nymphs, and fed nymphs, demonstrating partial blockade of *Bb* acquisition and transmission. Larvae from vaccinated mice (PBL+rCOM-M6) exhibited very low, mostly trace levels of *Bb* in the flat nymph stage, insufficient to transmit the pathogen to naïve mice. Larvae from vaccinated mouse (PBL+rCOM-M13) had undetectable *Bb* burden at all stages (Fig 8C).

These observations are consistent with seroconversion, where naïve mice challenged with nymphs derived from M6 or M13 did not seroconvert, whereas naïve mice challenged with nymphs from M2 developed strong *Bb*-specific antibody responses. These results indicate that a single oral PBL+rCOM vaccination effectively reduces or blocks *Bb* burden throughout larval, nymphal, and subsequent larval stages, with M4 showing partial reduction and M6 exhibiting minimal residual *Bb* insufficient for transmission. For this longitudinal tracking, at least 20 nymphs were required to perform all these analyses throughout all stages which limited the number of mice tracked in this experiment.

### Effects of oral vaccination with PBL+rCOM in *P. leucopus* hosts

The efficacy of the PBL+rCOM administered orally was also evaluated in *P. leucopus (Pl)* hosts, as they serve as the most common reservoir host for the Lyme disease agent in many parts of the US. It was also critical to connect the nature of the immune response induced by PBL+rCOM in *P. leucopus,* as this mammalian host is evolutionarily distinct from murine hosts and have immunological and physiological variations that need to be evaluated for modifying the dose and frequency of immunizations optimal for blocking *Bb* transmission from host to naïve *Is* larvae. *Pl* hosts are generally considered as immune-docile, with several parameters of the immune response that can be measured being homeostatic compared to murine hosts infected or immunized with pathogens/formulations, respectively (23, 24, 70).

Three groups (*n*=3) of 4-6 weeks old *P. leucopus* were vaccinated using a three-dose immunization regimen either via intradermal or via oral gavage, while unvaccinated animals were maintained as controls (Fig 9A, B). All animals immunized via intradermal or oral route had significantly higher antibody titers compared to unvaccinated controls at day 14 and 28 days post-vaccination (Fig 9D, E). All *Pl* hosts were challenged at day 29 by needle inoculation with 1x 10^5^ spirochetes per host with B31-AD strain in a volume of 100 uL intradermally (Fig 9C) and naive larvae, placed on day 56, were allowed to feed to repletion. qPCR analysis showed a significant reduction in the *Bb* burden in fed larvae from *Pl* hosts immunized with PBL+rCOM either via the intradermal or oral route compared to unvaccinated hosts (Fig 9F). The pathogen burden in the joint tissues was also significantly lower compared to that in unvaccinated hosts showing reduced pathogen persistence in tissues (Fig 9G). These observations provide a very strong premise for the utility of PBL+rCOM as an oral reservoir host targeted biologics to block transmission of *Bb* from infected *Pl* hosts to naïve *Is* larvae.

**Figure 9:**
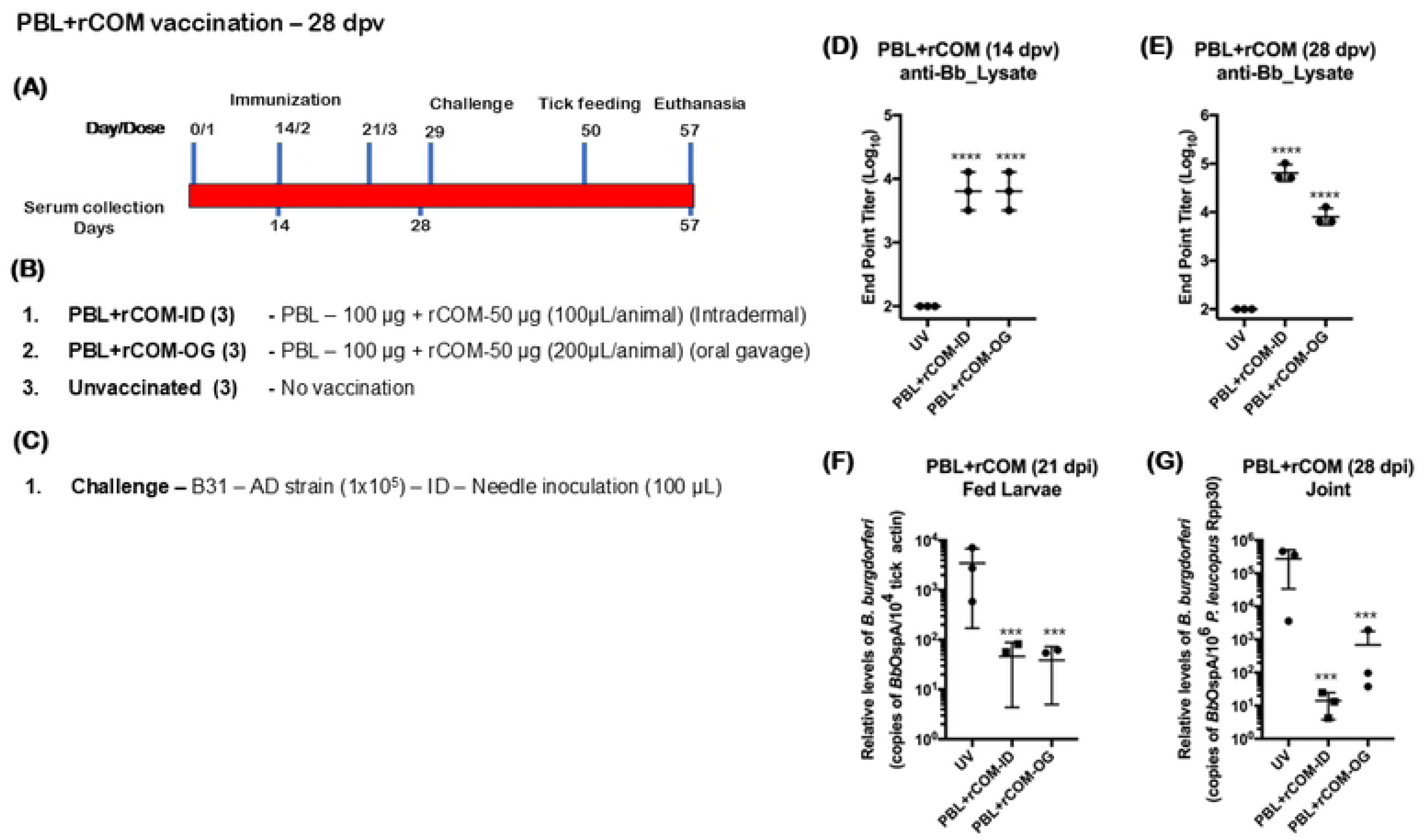
Efficacy of protection conferred by PBL+rCOM in *P. leucopus* against challenge with *B. burgdorferi* strain B31-AD strain. Schematic representation showing the timeline for immunization, challenge, euthanasia and serum collection time points (A), formulations, routes, and dosage used for immunization (B), and challenge strain and dose (C). Endpoint titers of serum antibodies from immunized mice at day 14 (D) or day 28 (E) post-vaccination against total *Bb* lysate determined by ELISA. Quantitative real time PCR analysis of spirochetal burden in skin in *Ixodes scapularis* larvae fed on challenged mice at day 21 post-infection with copies of borrelial *ospA* normalized to tick-actin copies (F). Quantitative real time PCR analysis of spirochetal burden in joints at day 28 post infection with copies of borrelial *ospA* normalized to *P. leucopus Rpp30* copies (G). Comparisons between vaccinated groups and unvaccinated controls were performed using a two-tailed Student’s *t*-test. Statistical significance is indicated as: *p < 0.05, **p < 0.01, ***p < 0.001, ****p < 0.0001.

### Effects of gender and challenge dose on efficacy of protection conferred by PBL+rCOM in *P. leucopus* hosts

We evaluated whether 1) the dose of the *Bb* inoculum administered via needle challenge and 2) gender influence the protective efficacy of PBL+rCOM in *P. leucopus*. As shown in Fig 10A and B, age- and sex-matched *P. leucopus* were vaccinated with a two-dose PBL+rCOM regimen administered via oral gavage. Seroconversion at day 14 was higher in male compared to female hosts, although variability was observed across vaccinated groups (Fig 10C), which is possibly due to differences in the oral response between animals. By day 28, antibody titers remained higher in males; however, variability persisted among both male and female vaccinated animals relative to naïve controls (Fig 10D). On day 35, animals were challenged intradermally with either 1x 10^3^ or 1x 10^5^ *Bb*-AD, and naïve *Ixodes scapularis* larvae were placed on day 56 and allowed to feed to repletion. qPCR analysis showed that *Bb* acquisition by larvae was significantly reduced in both male and female vaccinated hosts compared to unvaccinated controls (Fig 10E). As expected, *Bb* burden in fed larvae correlated with the challenge dose. Although antibody titers in *P. leucopus* were generally lower than those observed in similarly immunized C3H/HeN mice, there was a marked reduction in pathogen transmission to naïve larvae, highlighting differences in host immune responses between these models. Additionally, *Bb* burden in joint tissues was significantly lower in PBL+rCOM immunized *P. leucopus* compared to unvaccinated controls (Fig 10F), demonstrating that the formulation confers protective efficacy in both male and female reservoir hosts.

**Figure 10:**
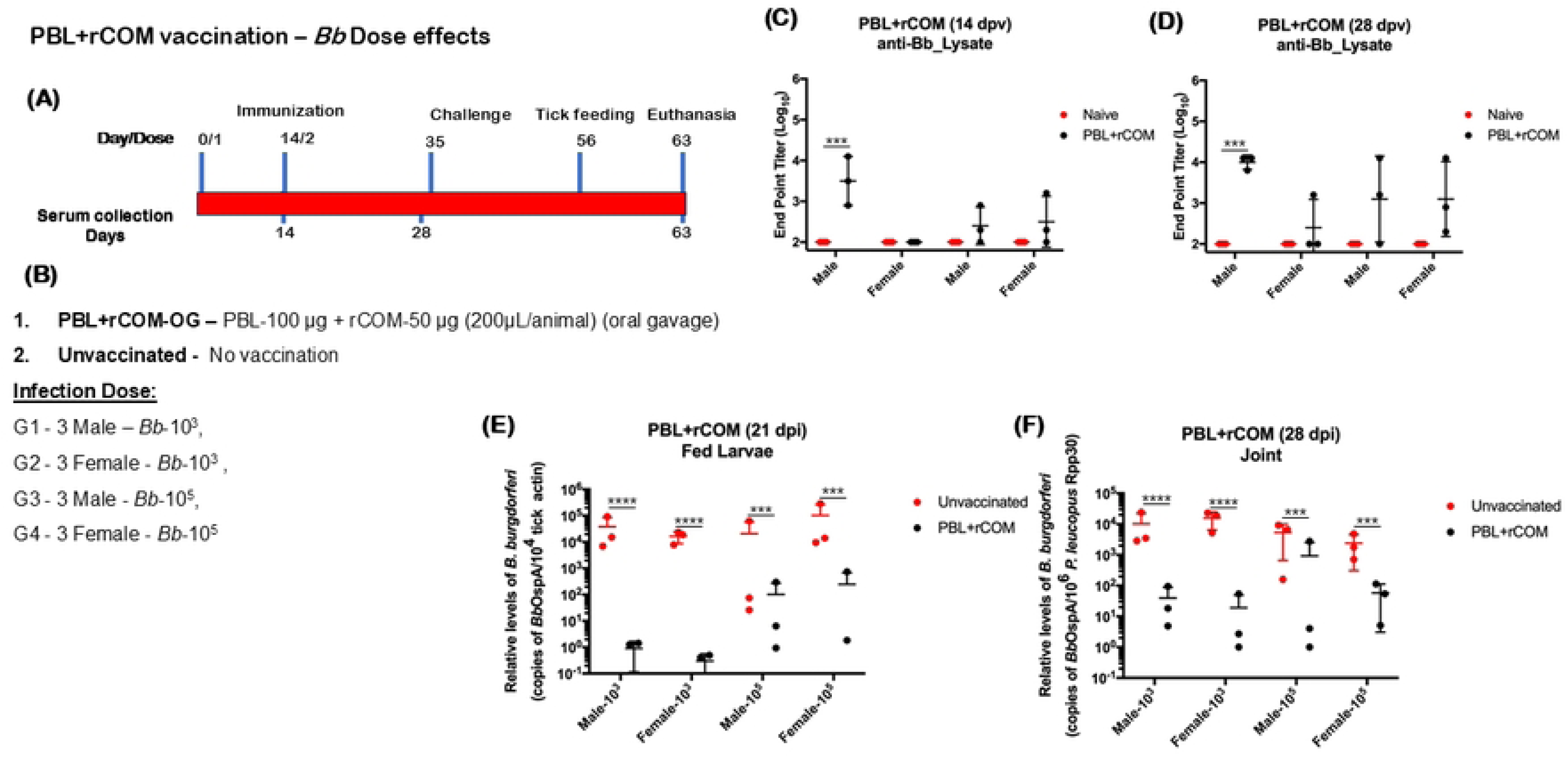
Efficacy of protection conferred by PBL+rCOM in *P. leucopus* against challenge with different doses of *B. burgdorferi* strain B31-AD strain. Schematic representation showing the timeline for immunization, challenge, euthanasia and serum collection time points (A) and formulations, routes dosage used for immunization and *Bb* challenge: 1x10^3^ and 1x10^5^ (B). Endpoint titers of serum antibodies from immunized mice at day 14 (C) or day 28 (D) post-vaccination against total *Bb* lysate determined by ELISA. Quantitative real time PCR analysis of spirochetal burden in *Ixodes scapularis* larvae fed on challenged mice at day 21 post-infection with copies of borrelial *ospA* normalized to tick actin copies (E). Quantitative real time PCR analysis of spirochetal burden in joints at day 28 post-infection with copies of borrelial *ospA* normalized to *P. leucopus Rpp30* copies (F). Comparisons between vaccinated groups and unvaccinated controls were performed using a two-tailed Student’s *t*-test. Statistical significance is indicated as: *p < 0.05, **p < 0.01, ***p < 0.001, ****p < 0.0001.

### Effects of buffer composition on efficacy of protection of PBL+rCOM

Based on the significant protection conferred by PBL+rCOM following oral gavage in *P. leucopus*, we sought to refine this pathogen-derived biologic to improve stability and immunogenicity while minimizing components with limited contribution to protective responses. To assess the role of 300 mM sodium chloride in the original formulation, we compared two buffer systems: 1) phosphate-buffered saline (PBS, pH 7.4; 138 mM NaCl) and 2) high-salt buffer (HSB, pH 7.4; 300 mM NaCl) for preparing PBL+rCOM and evaluated their impact on host immune responses and protective efficacy (Fig 11A, B). Oral administration of PBL+rCOM formulated in HSB induced significantly higher antibody responses at days 14 and 28 post-vaccination compared to PBS, even after two immunizations (Fig 11C, D). All animals were subsequently challenged intradermally with 1x 10^5^ *Bb*-AD. Importantly, *P. leucopus* immunized with PBL+rCOM prepared in HSB showed a significant reduction in *Bb* acquisition by naïve *Ixodes scapularis* larvae compared to those receiving the PBS formulation, demonstrating a straightforward improvement in vaccine efficacy (Fig 11E).Consistent with these findings, *Bb* burden in joint tissues was also significantly lower in PBL+rCOM in HSB vaccinated hosts relative to unvaccinated controls (Fig 11F). Together, these results highlight the critical role of buffer composition in optimizing PBL+rCOM as an oral reservoir-targeted vaccine, enhancing immune responses, reducing *Bb* survival, and limiting transmission from natural reservoir hosts to naïve tick larvae, thereby disrupting the enzootic life cycle of *Bb*.

**Figure 11:**
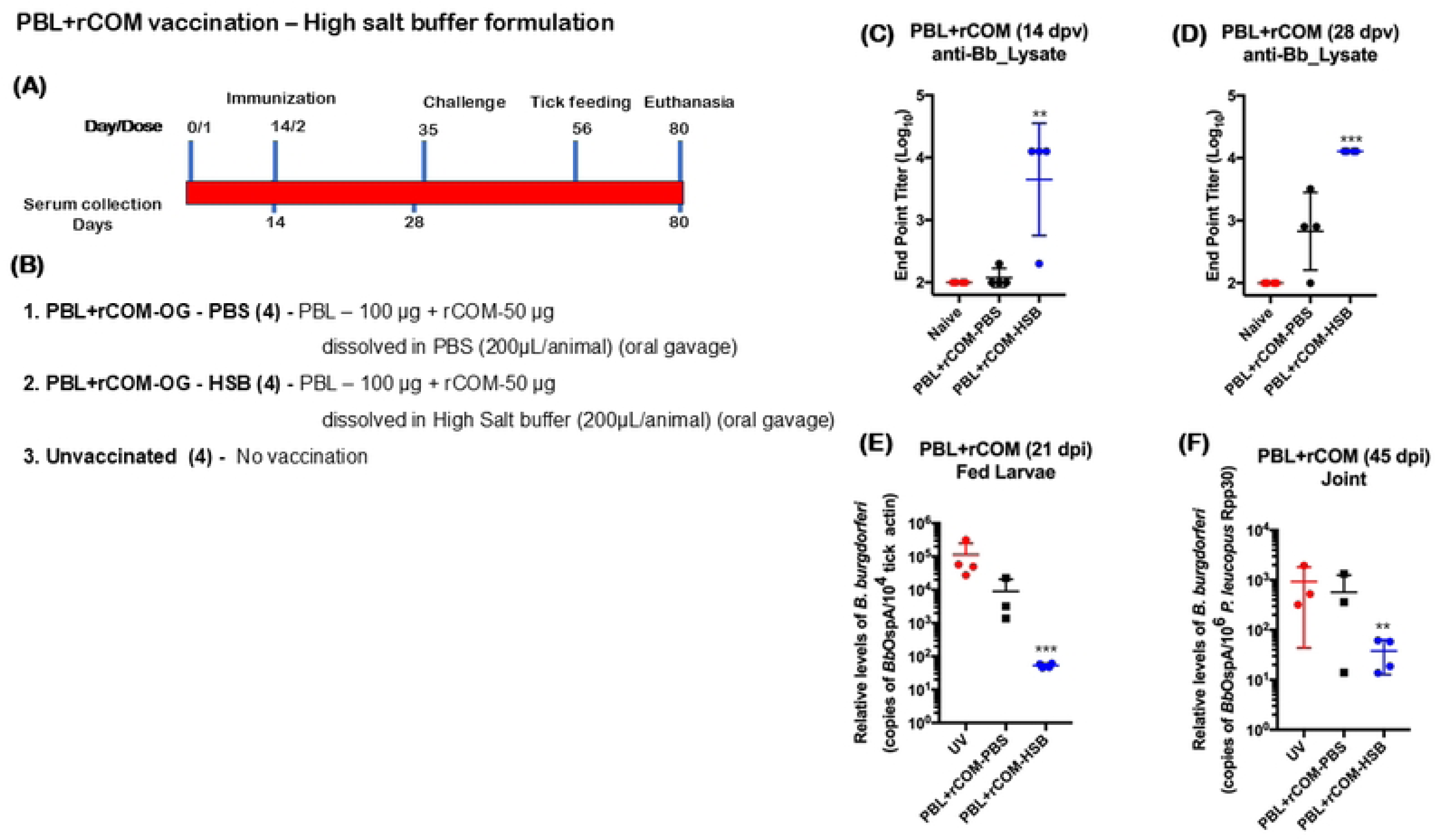
Efficacy of protection conferred by PBL+rCOM in high salt buffer in *P. leucopus* against challenge with *B. burgdorferi* strain B31. Schematic representation showing the timeline for immunization, challenge, euthanasia and serum collection time points (A) and formulations, routes dosage used for immunization and *Bb* challenge (B). Endpoint titers of serum antibodies from immunized mice at day 14 (C) or day 28 (D) post-vaccination against total *Bb* lysate, determined by ELISA. Quantitative real time PCR analysis of spirochetal burden in *Ixodes scapularis* larvae fed on challenged mice at day 21 post-infection with copies of borrelial *ospA* normalized to tick-actin copies (E). Quantitative real time PCR analysis of spirochetal burden in joint at day 45 post-infection with copies of borrelial *ospA* normalized to *P. leucopus Rpp30* copies (F). Comparisons between vaccinated groups and unvaccinated controls were performed using a two-tailed Student’s *t*-test. Statistical significance is indicated as: *p < 0.05, **p < 0.01, ***p < 0.001, ****p < 0.0001.

### High salt diet had no impact on PBL+rCOM induced protective immune response

Our findings indicated that increased sodium chloride is important for inducing immune responses in *P. leucopus*; however, sustained exposure to a high-salt diet could potentially alter vaccine-induced protection. To assess this, *P. leucopus* (three males and three females) were maintained on a high-sodium diet (4% NaCl) for two weeks, while animals on a standard diet (0.4% NaCl) served as controls. Both groups were then vaccinated with a two-dose PBL+rCOM regimen. The magnitude of antibody responses was comparable between animals maintained on control versus high-salt diets (Fig 12C-F), however, consistent with prior observations, seroconversion was lower in females compared to males. Animals were subsequently challenged intradermally with 1x 10^5^ *Bb*-AD on day 35. qPCR analysis revealed a significant reduction in *Bb*-specific DNA in fed larvae from vaccinated mice in both diet groups compared to unvaccinated controls (Fig 12G, H), which was consistent with reduced *Bb* burden in joint tissues (Fig 12I, J).Importantly, the high-salt diet did not alter *Bb* burden in unvaccinated mice or affect larval acquisition, indicating that dietary sodium has minimal impact on *Bb* pathogenesis in *P. leucopus*. Together, these findings suggest that while elevated NaCl in the vaccine formulation enhances immunogenicity, a high-salt diet does not adversely affect the protective efficacy of PBL+rCOM vaccination.

**Figure 12:**
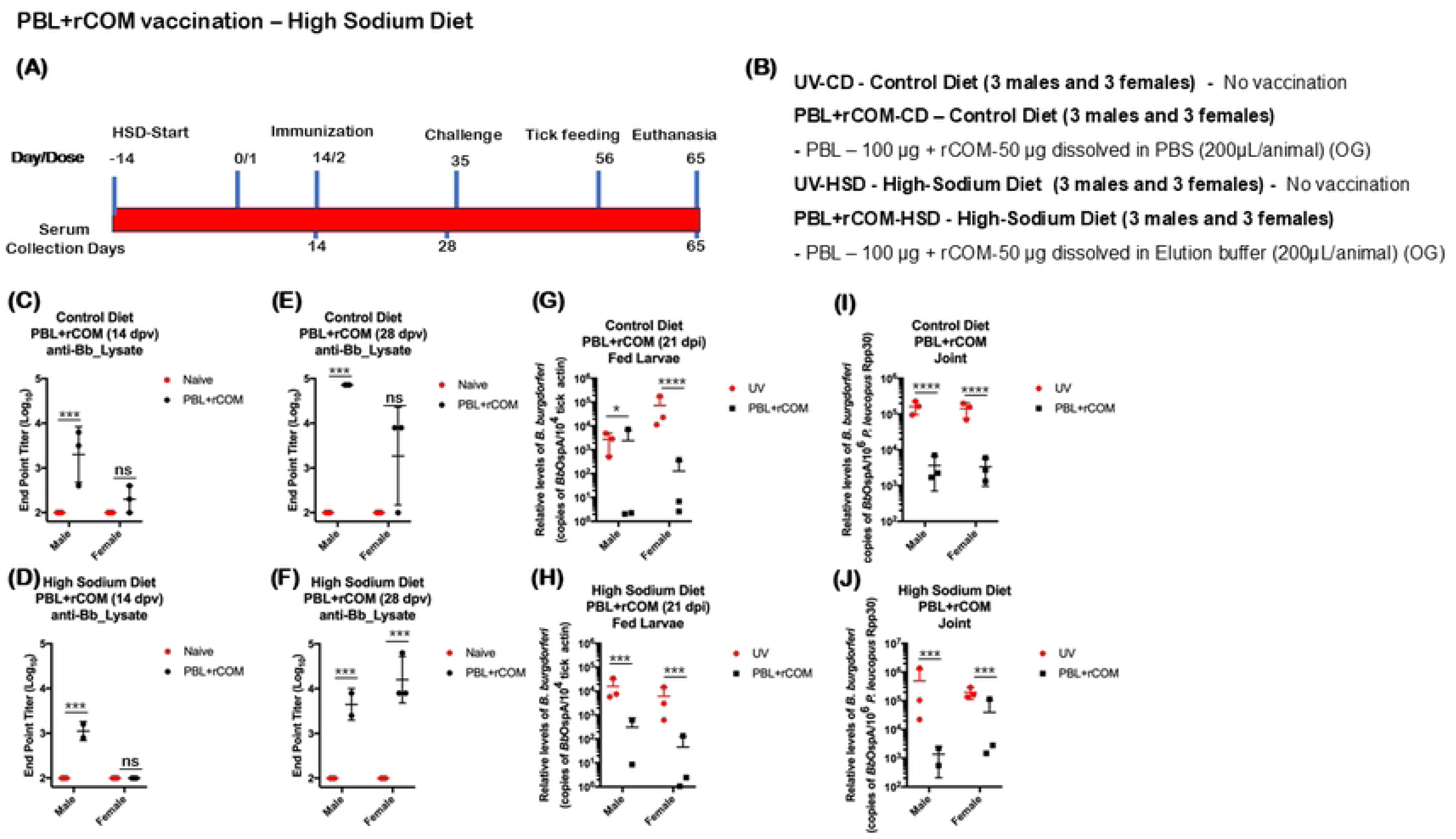
Efficacy of protection conferred by PBL+rCOM with high sodium diet in *P. leucopus* against challenge with *B. burgdorferi* strain B31. Schematic representation showing the timeline for feeding, immunization, challenge, euthanasia and serum collection time points (A) and formulations, routes dosage used for immunization and *Bb* challenge (B). Endpoint titers of serum antibodies from immunized mice on control diet or high salt diet at day 14 (C, D) or day 28 post-vaccination (E, F) against total *Bb* lysate determined by ELISA. Quantitative real time PCR analysis of spirochetal burden in *Ixodes scapularis* larvae fed on challenged mice at day 21 post-infection with copies of borrelial *OspA* normalized to tick-actin copies (G, H). Quantitative real time PCR analysis of spirochetal burden in joints at day 30 post-infection with copies of borrelial *OspA* normalized to *P. leucopus Rpp30* copies (I, J). Comparisons between vaccinated groups and unvaccinated controls were performed using a two-tailed Student’s *t*-test. Statistical significance is indicated as: *p < 0.05, **p < 0.01, ***p < 0.001, ****p < 0.0001.

### Efficacy of PBL+rCTB on short- and long-term protection in WFM

To further dissect the role of rCOM components in *P. leucopus*, we analyzed if rCTB alone added exogenously along with PBL will confer similar protective responses in the short-(Fig S4) and long-term (Fig S5) in *Pl* hosts and if the levels of rCTB can be varied to alter the host responses. As shown in Fig S4A and B, a single dose of PBL with rCTB given via the intradermal or oral gavage did induce significantly higher antibody responses at day 14 and 28 (Fig S4 D-G) compared to unvaccinated *Pl* hosts. Antibody titers measured 14 days (day 49) after needle challenge with B31-AD at day 35 showed significantly higher antibody responses in all vaccinated hosts compared to unvaccinated hosts and in unvaccinated challenged hosts (Fig S4H, I). Reduction in *Bb* burden in fed larvae at day 56 (21 days post-challenge) was not statistically significant between vaccinated and unvaccinated hosts; however, fed larvae from three of four intradermally (PBL+rCTB-ID) vaccinated hosts and three of five oral gavage (PBL+rCTB-OG) vaccinated hosts showed reduced burden, suggesting variable protection. (Fig S4J).

We repeated this study to determine the long-term protective responses with a single dose of PBL+rCTB (Fig S5A, B) and determined that the antibody responses were significantly higher at day 14 and day 174 post-vaccination with either *Bb* lysate or PBL used as the antigens in the ELISA based method used for determining the titers (Fig S5D-F). Furthermore, immunoblot analysis also revealed significant reactivity with several borrelial antigens as a part of the *Bb* lysate or PBL with serum collected at day 191 (14 days post-needle challenge intradermally, Fig S5G). There were also significantly higher antibody responses in the vaccinated challenged *Pl* hosts, as expected, compared to unvaccinated challenged or naïve hosts (Fig S5H, I). In addition, there was a significant reduction (p<0.05 *; p<0.01 **) in the levels of *Bb* burden in *Is* larvae allowed to feed on vaccinated *Pl* hosts compared to unvaccinated hosts (Fig S5J). These studies demonstrate that PBL+rCTB reduces the ability of naïve *Is* larvae to acquire *Bb*; however, the combination of PBL+rCOM (which also encodes for rCTB along with OspA+M cell-binding peptide) appears to be a more robust reservoir host-targeted oral formulation.

### Efficacy of protection of C3H/HeN mice and *Pl* hosts conferred by PBL+rCOM against *Bb* infected nymph challenge

A key criterion for validating PBL+rCOM as a reservoir-targeted oral formulation is its protective efficacy against *Bb* transmission via infected nymphs, to replicate the natural cycle, as the salivary components associated with tick feeding can modulate host immune responses. We therefore evaluated the efficacy of two doses of PBL+rCOM administered either intradermally or via oral gavage in both C3H/HeN mice and *P. leucopus* hosts (Fig 13A, B). Vaccinated animals showed significantly higher antibody titers at day 48 post-vaccination following oral administration of PBL+rCOM (Fig 13C). All animals were subsequently challenged on day 48 with *Bb*-infected *Is* nymphs (*n=*5 per animal), generated from larvae previously fed on a single needle-infected C3H/HeN mouse or *P. leucopus* host. ELISA using aqueous phase antigens (proteins that are not present in the PBL) revealed no significant differences between vaccinated and unvaccinated groups, confirming successful infection in all animals (Fig 13D). Consistent with this, qPCR analysis detected *Bb*-specific DNA in all fed nymphs (Fig 13E). However, *Bb* burden in nymphs feeding on vaccinated hosts was significantly reduced compared to those feeding on unvaccinated controls (C3H/HeN, p<0.05; *P. leucopus*, p<0.0001; Fig 13E), indicating that vaccine-induced antibodies reduced pathogen levels within feeding ticks. Furthermore, naïve larvae feeding on challenged vaccinated hosts exhibited lower *Bb* burden compared to those feeding on unvaccinated controls (Fig 13F).

**Figure 13:**
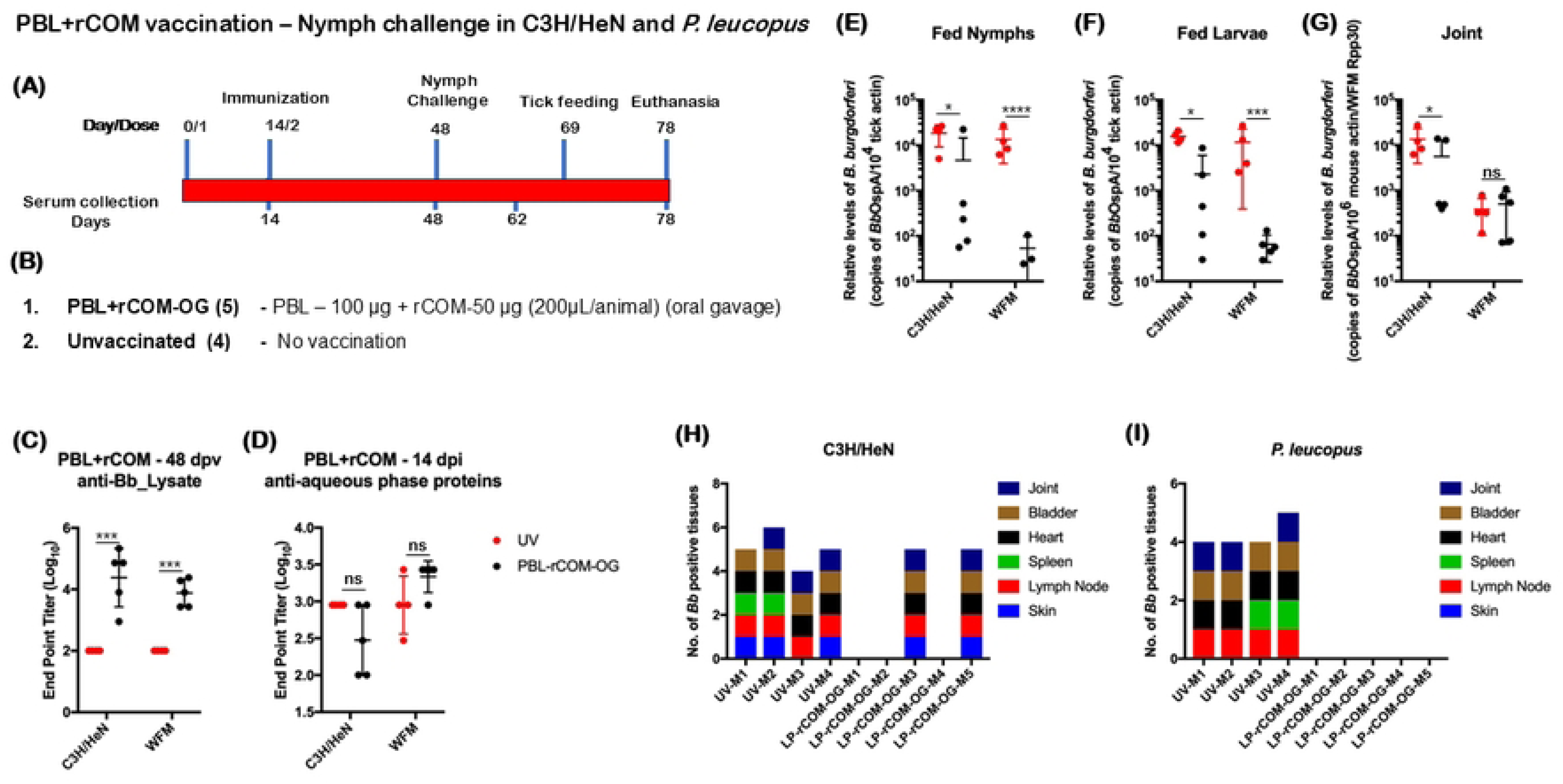
Efficacy of protection conferred by PBL+rCOM against nymph challenge with *B. burgdorferi* strain B31 in C3H/HeN mice and *P. leucopus*. Schematic representation showing the timeline for feeding, immunization, challenge, euthanasia and serum collection time points in C3H/HeN mice and *P. leucopus* (A) and formulations, routes, and dosage used for immunization and *Bb* challenge (B). Endpoint titers of serum antibodies against *Bb* lysate at 48 dpv (C) and serum antibodies from 14 days post-nymph challenge against aqueous phase protein (does not contain antigens from PBL) (D), determined by ELISA. Quantitative real time PCR analysis of spirochetal burden in *Bb* infected *Ixodes scapularis* nymphs after feeding to repletion on vaccinated and unvaccinated animals (E) and *Bb* burden in fed larvae at day 21 post-infection with copies of borrelial *OspA* normalized to tick-actin copies (F). Quantitative real time PCR analysis of spirochetal burden in joints at day 30 post-infection with copies of borrelial *OspA* normalized to mouse b-actin and *P. leucopus Rpp30* copies (G). Stacked column graph showing the recovery of viable *Bb* in six tissues from C3H/HeN mice (H) and *P. leucopus* (I) on euthanasia at day 78. Comparisons between vaccinated groups and unvaccinated controls were performed using a two-tailed Student’s *t*-test. Statistical significance is indicated as: *p < 0.05, **p < 0.01, ***p < 0.001, ****p < 0.0001.

In host tissues, *Bb* burden in joints was significantly reduced in vaccinated C3H/HeN mice, whereas no significant difference were observed in *P. leucopus*, although overall burden was lower than in mice (Fig 13G). Viable spirochetes were recovered from tissues in only two of five vaccinated C3H/HeN mice, corresponding to lower antibody responses in those animals, while no viable spirochetes were detected in any vaccinated *P. leucopus* hosts (Fig 13H, I). In contrast, all unvaccinated C3H/HeN and *P. leucopus* hosts had multiple tissues positive for spirochete growth in BSKII medium. Together, these findings demonstrate that oral administration of PBL+rCOM confers significant protection against *Bb*-infected nymph challenge and effectively reduces pathogen transmission from infected mammalian hosts to naïve *Ixodes* larvae, thereby supporting its potential as a reservoir-host targeted bait formulation to disrupt the natural transmission cycle of *Bb*.

## Discussion

The rapid rise in Lyme disease incidence across endemic regions of the United States, the widespread distribution of the competent vector *Ixodes scapularis* (*Is*) across nearly half of U.S. counties, the presence of multiple vertebrate reservoir hosts, the lack of a licensed human vaccine, and variable outcomes with antibiotic therapy collectively underscore the need for a multipronged and sustainable control strategy (71). This possible alternative strategy is to reduce the risk of acquiring Lyme disease. Reservoir host-targeted vaccination, particularly via oral delivery, represents a promising approach to interrupt the enzootic transmission cycle of *Bb*, thereby reducing the human incidence of Lyme disease. In this study, we systematically developed and evaluated Purified Borrelial Lipoproteins (PBL) as a pathogen-derived oral biologic, leveraging their native antigenic composition and compatibility with mucosal delivery platforms. We demonstrated the PBL-induced protective immune correlates in two important models: 1) C3H/HeN mouse model and 2) *P. leucopus* reservoir host model that serves as the major reservoir host that transmits *Bb* to larval ticks (22, 72).

PBL comprises a broad spectrum of immunogenic lipoproteins, including OspA, OspB, OspC, OspD, and BBK32, enriched via Triton X-114 detergent extraction that simultaneously inactivates the pathogen while preserving antigenic integrity. These lipoproteins are well-characterized modulators of host immune responses and can be potentially used to induce protective immune responses(73). Recently we demonstrated the immunogenicity following intradermal administration of PBL from *Bb* B31-A3 and two lipoprotein hyper expression mutants (delta *badR* and 8S strain) in C3H/HeN mice (45). Consistently, we observed strong reactivity of several antigens in PBL to infection-derived sera, providing a rationale for advancing PBL as a reservoir-targeted vaccine platform (Fig 1). PBL, comprised of enriched surface expressed antigens, may induce immune responses that target both host- and tick-phase *Bb* proteins, providing a unique strategy to disrupt the tick-pathogen-reservoir axis of Lyme disease (74).

Our initial studies demonstrate that oral administration of PBL-alone (PBL-OG), or as nanoparticle-encapsulated formulations (PBL-NP-OG), elicited significant antibody responses and reduced *Bb* burden in host tissues and feeding larvae (Fig 2). These findings establish proof-of-concept for oral delivery; however, follow-up studies revealed that while vaccination-induced seroconversion remained robust, residual spirochetes persisted in several tissues at later time points (60 dpi), suggesting partial waning of protective immunity (Fig S1). Notably, delivery of PBL as encapsulated nanoparticle using chitosan-PEG-PLGA did not substantially enhance durability, indicating that antigen composition alone may be insufficient to sustain long-term mucosal immunity (Fig 2 and Fig S1) (75). These findings suggest that PBL alone are sufficient to induce antibody responses without the need for a delivery system; however, a robust adjuvant is required to achieve durable, long-term protective immunity.

To address this, we explored the ability of mucosal adjuvants using recombinant OspA constructs with M-cell binding peptides (rOM), targeting the microfold cells in the gut-associated lymphoid tissue (GALT) administered via the oral route (Fig S2A) (69). Although rOM formulations with additional adjuvants such as Cholera Toxin B (CTB) and Eudragit enhanced antibody responses against total *Bb* lysate (Fig S3), they failed to reduce *Bb* burden or transmission, highlighting the immunogenicity of the lipid components of OspA and the need for an array of lipoproteins in their native conformations that comprise PBL (46). Since rOM with addition of rCTB induced better seroconversion, as CTB is a potent systemic mucosal adjuvant, we developed a modified construct, rCOM, incorporating CTB at the N-terminus followed by OspA and two repeats of M-cell targeting motifs (Fig S2B), which was used as an adjuvant in combination with PBL, throughout the study (76–78).

Iterative studies with PBL+rCOM revealed clear immune correlates of protection. Animals with higher antibody titers exhibited reduced *Bb* burden in tissues and diminished transmission to naïve larvae, whereas low responders retained both tissue infection and transmission capability (Fig 3). Also, the PBL+rCOM-OG vaccinated mice with significantly higher antibody responses, showed no spirochetes in the tissues and less transmission to tick larvae (Fig 3C, D), comparable to intradermal vaccinations. Importantly, rCOM alone was insufficient to confer protection, emphasizing the necessity of the multivalent antigenic composition provided by PBL. Alternative formulations, such as PBL with ragweed pollen grains (PBL-RPG-OG), also induced strong antibody responses, further supporting the flexibility of the PBL platform for oral delivery. This adaptability enables the incorporation of diverse delivery systems, including nanoparticles and pollen grains, as needed to protect gut acid-sensitive protein antigens and enhance formulation stability (65).

Immunization using two-dose regimens demonstrated that both intradermal and oral administration of PBL+rCOM induced significant antibody responses and reduced *Bb* acquisition by naïve larvae (Fig 4). Also, there are no viable spirochetes recovered from the tissues of the vaccinated mice. In the previous experiment (Fig 3), even after 3 doses of PBL+rCOM-OG, one out of four animals did not induce significantly higher response and that was reflected in all readouts such as *Bb* burden in skin and joint as well as acquisition by naïve larvae, suggesting significant antibody responses are the key immune parameters for oral vaccination. It is to be noted that PBL-OG induced antibody responses were significantly lower than PBL+rCOM-OG at 74 dpv, suggesting that rCOM is critical for consistent antibody responses (Fig 3C). Overall, these two experiments (Figs 3 and 4) demonstrated that PBL+rCOM-OG consistently met three key criteria of a potential efficacious oral vaccine: 1) robust peripheral immune responses, 2) reduced transmission of *Bb* to larvae and 3) reduced *Bb* burden in tissues.

Our study using PBL is directed at expanding the antigenic repertoire of *Bb* to drive prolonged and robust immune responses adding to the protective efficacy of OspA-based reservoir host-targeted formulations using bacterial or viral vectors that that have been validated in several prior studies (15, 17–19, 27, 40, 44). Several other formulations comprising of antigens from *Bb* that are a part of PBL have also been developed and tested for efficacy of protection in mouse models of Lyme disease with the intention to advance these formulations as human vaccines (42, 43, 79–86). Although recombinant OspC expressed in *E. coli* delivered as an oral vaccine did not protect mice from *Bb* challenge via ticks presumably due to multiple phyletic types of *Bb* OspC (87), our formulation had OspC at stoichiometric levels that reflect the composition of all major borrelial lipoproteins on the surface of *Bb*. A significant advantage of using PBL as oral biologics is to expand the antigenic repertoire of *Bb* that broaden the immune responses against a variety of immunogenic lipoproteins to target *Bb* both within the ticks (via anti-OspA antibodies) as well as during colonization of various host tissues (via anti-OspC and other surface exposed lipoprotein specific antibodies). Overall, our study was based on multiple check points to efficiently interfere with tick-pathogen-reservoir host axis of Lyme disease (15, 16, 86, 88)

Comprehensive immunological analyses further demonstrated that PBL+rCOM induced broad humoral responses, including elevated total IgG, multiple IgG isotypes, and IgM, along with mucosal IgA responses in fecal samples (Fig 5). In parallel, antigen-specific T cell responses, characterized by IFN-γ and IL-4 production, were significantly increased at 28 and 65 dpv, suggesting the induction of both Th1 and Th2 memory responses (Fig 6). While IgG1 and Th2 responses are potentially through incorporation of CTB as part of the formulation (89), whereas IgG2b and Th1 responses may be driven by enhanced antigen uptake via M-cells together with the intrinsic immunostimulatory properties of PBL (45, 63). These cellular responses likely contribute to the durability of protection, as evidenced by long-term studies showing sustained antibody responses (Fig 7B, D) and transmission-blocking efficacy for up to 10 months following a single oral dose (Fig 7E).

Critically, evaluation of the full enzootic cycle demonstrated that vaccination reduced *Bb* burden, not only in hosts, but also across tick developmental stages. Notably, the nymphs molted from larvae fed on vaccinated, *Bb* infected mice did not transmit spirochetes to naïve animals (Fig 4I, J). Additionally, nymphs collected from long-term vaccinated mice also showed reduced transmission to naïve animals (Fig 8). Although sterilizing immunity was not uniformly achieved, there was a marked reduction in transmission from infected nymphs derived from vaccinated hosts. These findings reinforce the concept that reservoir-targeted interventions need not achieve complete pathogen clearance but rather reduce pathogen prevalence below transmission thresholds, sufficient to disrupt the natural infection cycle (90).

Translation to the natural reservoir host, *P. leucopus*, further validated the efficacy of PBL-based formulations. We used *Bb*-AD strain for infection studies in *P. leucopus*. A three-dose regimen of PBL+rCOM (Fig 9) via intradermal and oral routes elicited robust antibody responses and significant reductions in *Bb* transmission to naïve larvae, consistent with observations from C3H/HeN mice. Also, PBL+rCOM-OG induced protection was consistent across sexes (Fig 10), where higher challenge doses (1x10^5^) of spirochetes revealed dose-dependent variability, while lower doses (1x10^3^) are completely cleared. Although higher challenge dose exhibited more *Bb* burden in tissues and larvae, vaccination consistently reduced *Bb* burden in joint tissues and transmission to larvae in both doses compared to the unvaccinated controls. This supports that PBL+rCOM-OG significantly reduces even higher doses of *Bb* challenge, consistent with previous findings of reduced *Bb* colonization in ticks in a dose-dependent manner with the circulating anti-OspA IgG concentration (91). To evaluate the necessity of rCOM in *P. leucopus*, we tested whether PBL with recombinant CTB as an adjuvant could elicit protective immunity. Although single dose of PBL+rCTB-OG induced significantly higher antibody responses at day 14 and 174 post vaccination via either intradermal or oral route, it either failed to reduce transmission of *Bb* to larvae at 56 days post-vaccination (Fig S4) or was not efficacious at blocking transmission at 198 days post-vaccination (Fig S5), underscoring the requirement for rCOM as an adjuvant for PBL-based formulation for *P. leucopus*.

It is important to factor significant evolutionary differences in the physiological and immunological parameters of protection between C3H/HeN and *P. leucopus* hosts (67, 90, 92, 93). While a variety of reagents are available to characterize immune responses in C3H/HeN mice, our ability to interrogate responses to pathogen-derived biologics, adjuvants, and infection-induced immunity in *P. leucopus* is limited by the lack of species-specific reagents. Consequently, our correlates of protection are largely restricted to IgG levels and reductions in pathogen transmission and prevalence of *Bb* in ticks and host tissues, respectively (94–96). Despite these limitations, we observed comparable levels of protection following immunization of PBL+rCOM via intradermal and oral routes between C3H/HeN and *P. leucopus* hosts reflecting the potential for use of these formulations across hosts. There is a dire need to advance these reservoir-host-targeted formulations to confer protection in a wide array of vertebrates that serve as reservoirs of tick-borne pathogens as well as develop platforms to incorporate antigens from a multiple tick-borne pathogens that are transmitted as single or coinfections (97).

Another important component of PBL+rCOM formulation is sodium chloride (NaCl). Specifically, NaCl concentration influenced both immune responses and transmission outcomes (Fig 11). Formulation of PBL+rCOM in a buffer containing 300 mM NaCl (pH 7.4) significantly enhanced antibody responses and reduced transmission compared to formulations prepared in 120 mM NaCl (pH 7.4). Several studies have demonstrated that NaCl can modulate gut microbiomes, which in turn influences immune pathways, including the expansion of Th17 cells and the development of B cell responses (98–100), warranting further mechanistic investigation in this context. Given that the intended field application involves delivery via feed pellets, we further evaluated whether a high-sodium diet alters PBL-induced immune responses. *P. leucopus* maintained on control versus high-sodium diets showed no significant differences in PBL+rCOM-OG induced immune correlates (Fig 12). These findings suggest that incorporation of higher sodium content into feed pellet formulations is feasible and does not significantly compromise the vaccine-induced protective immune response.

Although most challenge studies were performed via needle inoculation to maintain a uniform infectious dose, optimize oral vaccine formulations, and enable direct comparisons across groups (Fig 1-12); tick challenge experiments (Fig 13) confirmed that PBL+rCOM-OG remains effective under natural transmission conditions, where tick saliva and feeding-associated immunomodulatory factors play critical roles. In both vaccinated and unvaccinated C3H/HeN and *P. leucopus*, infection was established, as evidenced by *Bb*-specific antibody responses measured using aqueous phase antigens (distinct from PBL). However, *Bb* burden in feeding nymphs was significantly reduced in vaccinated animals, likely due to PBL-specific antibodies in the blood meal, particularly those targeting tick-phase *Bb* proteins such as OspA (42, 101). Notably, viable spirochetes were recovered from tissues of two C3H/HeN mice with lower antibody responses, but not from those with higher antibody levels, and were absent in all vaccinated *P. leucopus*. These findings further emphasize that vaccine-induced antibody responses are critical for reducing *Bb* burden in ticks, limiting host infection, and decreasing larval acquisition. This reduction in pathogen load translated into decreased transmission to subsequent developmental stages of the ticks underscoring the impact of vaccination in interfering with the life cycle of *Bb*.

In summary, this study establishes PBL+rCOM as a robust, pathogen-derived, oral reservoir host-targeted biologic that fulfills two critical criteria: 1) induction of long-term protective immune responses and 2) significant reduction in *Bb* transmission to naïve larvae. While challenges remain, including optimization of formulation stability and delivery for field deployment, this platform offers a flexible and scalable strategy for efficient blocking of the enzootic life cycle of *Bb.* Moreover, its adaptability to incorporate additional antigens provides a foundation for developing novel reservoir host-targeted vaccines targeting multiple tick-borne pathogens that are transmitted to humans as single or coinfections by *Is* ticks.

## Materials and Methods

### Animals and Ethics Statement

All animal experiments were performed in BSL2 facility within Laboratory Animal Resources Center (LARC), an AAALAC International Accredited facility at University of Texas-San Antonio. 6-8-week-old female C3H/HeN and BALB/c mice (Charles River Laboratories, Wilmington, MA) was used in this study. Breeding pairs for *P. leucopus* mice were obtained from the *Peromyscus* Genetic Stock Center at University of South Carolina and were bred following the harem breeding protocol approved at UTSA. All animal experiments were conducted following USDA guidelines for housing and care of laboratory animals and in accordance with protocols approved by the Institutional Animal Care and Use Committee of UTSA. Based on these guidelines, general condition and behavior of the animals were monitored by trained laboratory and LARC staff daily and methods to minimize pain and discomfort were adopted as needed in this study. All tick infection studies were carried out within environmental chambers located within the BSL2 facility within LARC at UTSA. All animals were maintained on normal diet, while specific groups were maintained on a high sodium diet (4% NaCl grain diet, TestDiet).

### Bacterial strain and Growth conditions

A low passage infectious clonal isolate of *Bb* strain B31-A3, kindly provided by Patricia Rosa (Rocky Mountain Laboratories, NIAID, NIH, Hamilton, MT) (102) and *Bb* strain B31-AD, adapted in *P. leucopus*, generously provided by Jeffrey Bourgeois and Linden Hu (Tufts University, Boston, MA) (103) were propagated at 32°C or 37°C in liquid Barbour-Stoenner-Kelly (BSKII, pH 7.6) media supplemented with 6% heat inactivated rabbit serum (Pel-Freez Biologicals, Rogers, AR) as previously described (104–109). Once the cultures reached a density between 1-2 x 10^7^ spirochetes/mL, viable spirochetes were enumerated by dark field microscopy and used for infection studies after ensuring presence of all critical infection associated plasmids by PCR needed for spirochete survival in ticks and mammalian hosts.

### Lipoprotein purification from *Bb*

Lipoproteins were extracted from the spirochetes as previously described (44, 45). Briefly, 1x10^9^ *Bb* was solubilized in 1 mL of PBS (Phosphate Buffered Saline) pH 7.4 containing 1% Triton^®^ X-114 (TX-114) (Sigma Aldrich) by gentle rocking at 4°C overnight. Pellet containing the TX-114 insoluble material was removed by centrifugation twice at 15,000 x *g* at 4°C for 15 mins. The supernatant was transferred to a sterile tube and incubated at 37°C for 15 mins followed by centrifugation at 15,000 x *g* for 15 mins at RT. The aqueous phase was transferred to a new tube and re-extracted one more time with 1% TX-114 as described above. The detergent phase was washed with 1 mL PBS pH 7.4 thrice and the detergent phase proteins were precipitated by adding 10-fold volume of ice-cold acetone. Precipitates were collected by centrifugation at 15,000 x *g* at 4°C for 30 mins, dried to remove acetone, and stored at -20°C until further use. Insoluble materials (Pellet), soluble proteins (aqueous phase) and Lipoproteins (TX-114 phase) were analyzed by SDS-PAGE gel (Fig 1). Purified Borrelial Lipoproteins (PBL) were quantified using a BCA assay kit (Thermo Scientific).

### Design, construction, overexpression and purification of recombinant multivalent antigen

To improve the efficacy of the vaccine, we synthesized two gene fragments (Twist Bioscience), where rCTB-OspA-Mcell (rCOM) with a 6X-Histidine tag at the N-terminus to purify the recombinant protein, TEV protease site to remove the His-tag, Cholera Toxin B as an oral vaccine adjuvant, borrelial Outer surface protein A (OspA) as the protective antigen, and two repeats of M-cell peptide sequence targeting the microfold cells present on the Peyer’s patches at the C-terminal end connected by linker sequences. while rOspA-M-cell comprised a similar construct except Cholera Toxin B. The constructed gene fragments were cloned into a pET23a plasmid, transformed into electrocompetent TOP10 *Escherichia coli* (Invitrogen, Carlsbad, CA). Gene sequences and the primers used are listed in the Supplementary Table 1. Positive colonies containing the amplified gene were picked to purify the cloned expression vector, confirmed by sequencing using T7 promoter and T7 terminator primers. Plasmids verified by sequencing were transformed into the *E*. *coli* expression host, Rosetta (Novagen) and expression was induced with 1 mM IPTG for 4 hours at 37°C in a 500 mL flask and the culture was pelleted by centrifugation for 10 min at 3000 x *g* (SORVALL, F13S rotor) followed by resuspension in Lysis buffer (50 mM NaH2PO4, 30 mM NaCl and 10 mM Imidazole, pH 8.0) containing a protease inhibitor cocktail (Pierce, Thermofisher). Resuspended cells were then lysed using a French press. The resulting pellet was solubilized in 1 mL of PBS (Phosphate Buffered Saline) pH7.4 containing 1% Triton^®^ X-114 (TX-114) (Sigma Aldrich) by gentle rocking at 4°C overnight. Pellet containing the TX-114 insoluble material contained the inclusion bodies of recombinant proteins and the pellet was washed twice with PBS and washed with ice-cold acetone. Precipitates were collected by centrifugation at 15,000 x *g* at 4°C for 30 mins, dried to remove acetone, and stored as pellets in -20°C. Recombinant proteins were analyzed on an SDS-PAGE gel for purity and quantified using BCA assay Kit (Thermo Scientific).

### Encapsulation of PBL

To improve the stability of the PBL in the gut and for sustained release of the proteins, we encapsulated the PBL with chitosan nanoparticles, ragweed pollen grains and Eudragit.

#### Chitosan encapsulation

In 3 mL of dichloromethane, 100 mg of PLGA polymer & 1 mg of *Bb*LP were dissolved. In 1% glacial acetic acid, 5 mL of chitosan solution (12% w/v) was made, filtered & then titrated to 5 mL of PVA aqueous solution (2% w/v) before emulsification. Thereafter, PEG (2 kDa), 2% w/v, was provided to the prepared chitosan-PVA aqueous solution prior to emulsification to gain PEG-coated (PLGA-CS-PEG) NPs. In their organic phase PLGA and PBL were emulsified in the previous mentioned chitosan-PEG-PVA aqueous solution, then sonicated by a microtip probe sonicator adjusted at energy output of 55 W in an ice bath for 2 min. The organic solvent was evaporated by stirring the emulsion at room temperature on a magnetic stirring plate overnight. In the next day, the emulsion was centrifuged at 21,000 x *g* at 4°C for 20 minutes was done to obtain the formed NPs, then the unbound PVA, chitosan, PEG and un-encapsulated PBL were removed by double distilled water washing. Finally, the encapsulated PBL were lyophilized and stored at -20°C until further use.

#### Ragweed Pollen encapsulation

Pollen grains of Ragweed, *Ambrosia elatior* were purchased from Pharmallegra, Czech Republic. Pollen grain shells were prepared as previously described. Briefly, 50 g of dry pollen grains were stirred in 300 mL of acetone under reflux overnight. Following filtration and overnight drying, they were stirred under reflux in 450 mL of 2 M potassium hydroxide for 12 h at 120 °C (renewed after 6 h), further filtered and washed with hot water (5 × 300 mL) and hot ethanol (5 × 300 mL). After drying overnight, dried pollen grains were stirred under reflux for 7 days in 450 mL of orthophosphoric acid at 180 °C. LS were filtered and washed sequentially with water (5 × 300 mL), acetone (300 mL), 2 M HCl (300 mL), 2 M NaOH (300 mL), water (5 × 300 mL), acetone (300 mL) and ethanol (300 mL) and dried at 60 °C overnight. Dry and processed ragweed pollens (0.2 g) were added to 2 mL aqueous lipoprotein solutions to obtain mixtures containing 20-100% w/w of lipoproteins with respect to pollens. The mixtures were incubated with gentle shaking overnight at 4°C, centrifuged at 3000 × g, and the supernatant was discarded to remove excess aqueous lipoprotein solution. Pellets were dissolved in 2X FSB, boiled for 5 minutes and were run on SDS-PAGE gel to check the presence of lipoproteins. Pellets with and without lipoprotein encapsulation were resuspended in double distilled water and boiled for 5 minutes and the supernatant was used to quantify the protein concentration using BCA assay. Pellets were resuspended in Saline solution (0.9% NaCl) to a final concentration of 100 µg/100 µL and used as the oral vaccine.

#### Eudragit encapsulation

To evaluate the efficacy of oral delivery of recombinant antigens, recombinant OspA fused with two tandem repeats of M cell (rOM) with or without the adjuvant recombinant cholera toxin B subunit (rCTB) were encapsulated using Eudragit L100, a polymer widely used for enteric protein delivery (110). Briefly, 250 mg of Eudragit L100 was dissolved in 10 mL of 100% ethanol and vortexed for 10 minutes until completely solubilized. Acetone-precipitated rOM (1 mg/mL) with or without rCTB (1 mg) was prepared in parallel. One milliliter of the antigen preparation was then added to 4 mL of the Eudragit solution and mixed thoroughly by vortexing for 1 minute. The mixture was subsequently vacuum-dried using a Vacufuge for 1 hour to remove ethanol. The resulting dried pellet was washed with saline (0.9% NaCl) and centrifuged at 15,000 × g for 15 minutes. The final pellet was stored at −20°C until further use. Pellets were dissolved in 100µL of 2X sample buffer (FSB), boiled for 5 minutes and were run on SDS-PAGE gel to check the presence of rOM and rCTB by Coomassie blue staining. To assess protein concentration, the pellet was resuspended in 100µL of sterile distilled water, and concentration was quantified using a BCA assay.

### Vaccination strategies

We conducted several vaccination experiments with different cohorts to check the protective efficacy of PBL with different combinations in mouse strains, C3H/HeN. For vaccination, we used 100 µg of PBL with or without nanoparticle/ragweed encapsulation in combination with or without 100 µg of rCOM dissolved in the buffer (50 mM NaH_2_PO4, 30 mM NaCl and 250 mM Imidazole, pH 8.0). In select experiments, PBL and rCOM were dissolved in 1X PBS. Aliquots of 100 µL of antigen were delivered via a ball tipped disposable feeding needle (Fisher Scientific, Pittsburgh, PA) for oral gavage inoculation on Day 0 and Day 14. For intradermal injections, 100 µL of formulation containing 50 µg of PBL and/or 50 µg of rCOM were injected into the ventral side using a 27-gauge needle.

### Serum and Feces collection for antibody titers

To determine the peripheral antibody levels, blood was collected from the lateral saphenous vein using Microvette CB 300 capillary tubes (Sarstedt AG & Co.KG), centrifuged at 4000 x *g* for 10 minutes and the supernatant were collected and stored at -20°C until further use. To determine the mucosal antibody levels in the gut, wet feces pellets were directly collected in sterile tubes, diluted in 1 mL PBS containing metabolic inhibitor, sodium azide (NaN_3_) and bovine calf serum as a protein carrier to reduce non-specific binding of antibodies and shaken gently overnight at 4°C. The extract was centrifuged twice at 4°C, 16,000 × *g* for 15 min. The supernatant was collected and stored at -20°C until further use.

### Enzyme Linked Immunosorbent Assay (ELISA)

All the assays were performed using a Costar High binding Assay Plate (Corning). ELISA was performed against borrelial lysate or PBL as required. 10 µg of the antigen were resuspended in 10 mL of coating carbonate buffer (0.05M, pH 9.6) and 100 µL of coating antigen were added to each well and incubated overnight at 4°C. After washing three times with Tris Buffered Saline supplemented with Tween 20 (TBST), 100 µL of 1% Bovine Serum Albumin (BSA) (Probumin, Millipore) in TBST was added to each well and incubated for 1 hr at room temperature and washed three times with TBST. Serum and feces samples were diluted in TBST starting from 1:50 up to 1:6400 dilutions were prepared and 100 µL of each diluted sample was added. After two hours of incubation at room temperature, plates were washed five times with TBST and 100 µL of 1:3000 diluted secondary anti-mouse total Ig, IgG, IgG1, IgG2a, IgG2b, IgG3, IgM and IgA antibodies (Southern Biotech) or anti-*P. leucopus* IgG antibodies (LGC Clinical Diagnostics) conjugated with Horseradish peroxidase (HRP) were added and incubated for 1 hr. After four washes, 100 µL of OPD substrate solution was added and read at 450 nm using Spark 10M (Tecan) microplate reader.

### Animal injections and tick larvae feeding

For infection studies, *Bb* cultures were verified by PCR for the presence of the virulence plasmids lp25 and lp28-1 on the day of injection. After confirmation, the cultures were centrifuged at 4000 x *g* for 20 minutes at 4°C and the pellets were washed three times with sterile HBSS at 4000 x *g* for 5 minutes at 4°C and the cells were resuspended in BSKII medium supplemented with 6% heat inactivated rabbit serum at a final concentration of 10^6^ spirochetes/mL. Each mouse was injected with 100 µL of spirochetes via intra-dermal injections. After 28 days of needle challenge, animals were moved to the tick facility, hairs were shaved on the dorsal side of the mouse and sterile plastic tubes were glued on to the skin of each mouse using special adhesive material. After overnight drying, 50-100 *Ixodes scapularis* larvae were loaded into the tubes and the tubes were capped to increase the tick movement towards the skin of each mouse. After 3 to 7 days, fed larvae were collected and stored in a sterile tube covered with perforated parafilm and incubated at 23°C under humid conditions for 6-8 weeks to allow the larvae to molt into nymphs. After collecting the fed larvae, the mice were moved to the regular BSL2 facility for necropsy. All mice infection studies were conducted in a Biosafety Level 2 facility.

### Nymph Challenge

To generate *Bb* infected nymphs for challenge studies, naïve larvae were allowed to feed on C3H/HeN mice that had been infected intradermally with 1×10^5^ *B. burgdorferi* strain B31-A3 as previously described (107). For *P. leucopus*, 1×10^5^ *B. burgdorferi* strain B31-A3-AD strain was used. To assess spirochete acquisition in flat nymphs, five nymphs from the stock tubes were crushed individually and analyzed for spirochete burden, as mentioned in the previous paragraph and reported as relative copy numbers of *Bb*FlaB to *I. scapularis* actin. After confirming the *Bb* load, infected nymphs from C3H/HeN and *P. leucopus* were then used to challenge naïve or immunized C3H/HeN mice and *P. leucopus* respectively (with or without vaccination) by allowing ticks to feed to repletion (3-7 days).

### Necropsy and quantitation of *Bb* burden in tissues

Mice were euthanized by carbon dioxide exposure followed by cardiac puncture, serving as terminal bleed and as secondary method of euthanasia. Skin, lymph nodes, spleen, heart, bladder and joint tissues were aseptically isolated and inoculated into BSKII medium and scored for growth, after a single blind pass at day 5, by examining cultures using dark field microscopy at day 14 post-isolation in primary and blind-pass cultures. Another set of skin, lymph nodes and spleen samples were stored at -80°C for genomic DNA extraction.

### Genomic DNA extraction and qPCR

Genomic DNA from the skin, spleen and lymph nodes from mice or *Ixodes scapularis* larvae and nymphs was extracted using High Pure PCR Template Preparation Kit (Roche, Indianapolis, IN). Briefly, samples were lysed with 200 µL of lysis buffer in a BeadBlaster™ 24 microtube homogenizer using Zirconia beads. Further, 40 µL of proteinase K (20 mg/mL) and 20 µL of Collagenase (1 mg/mL) was added and incubated overnight at 57°C. Next, DNA was extracted as directed by the manufacturer using the column. *Bb* burden in the samples was determined by a SYBR-based qPCR assay (PowerUp SYBR Master Mix, Applied Biosystems) using the *Bb*FlaB gene or *Bb*OspA gene specific primers for B31-AD strain and normalized to tick actin or mouse actin or *P. leucopus Rpp30* (111, 112) and the reaction was performed in a StepOne Plus real time PCR system (Applied Biosystems). Primers used are listed in Supplementary Table 1. Standard curves for all three genes were performed by full length genes encoded pCR2.1 plasmid and the results are presented as ratio of *Bb*FlaB or *Bb*OspA normalized to tick actin, mouse actin or *P. leucopus* Rpp30 copy numbers.

### ELISPOT

The frequency of PBL-specific T cells was analyzed by a standard IFN-γ and IL-4 ELISPOT assay (MabTech, Cat. # 3321-2 H). Splenocytes (5 × 10^5^ cells per well) were stimulated with 1 µg mL^−1^ of PBL. Cells treated with PMA (1 μg ml^-1^), Ionomycin (2.5 μg ml^-1^) were used as positive controls and PBS as negative controls. Spots were counted using the ELISPOT reader.

### Statistical Analysis

Statistical analysis of ELISA, ELISPOT and qPCR was determined using two-tailed unpaired Student’s t-test in GraphPad. Comparisons between vaccinated groups and unvaccinated controls were performed using a two-tailed Student’s *t*-test. Statistical significance is indicated as: *p < 0.05, **p < 0.01, ***p < 0.001, ****p < 0.0001 or as p-values represented in each graph.

## Acknowledgements

This study was partly supported by Public Health Service grant 5R01AI152233 from the National Institute of Allergy and Infectious Diseases (JS), pre-doctoral fellowship from the South Texas Center for Emerging Infectious Diseases (TMI). JFSL is supported by the UTSA Graduate School and The National Institute of Allergy and Infectious Diseases of the National Institutes of Health under Award Number T32AI184340. We sincerely thank Jeffrey Bourgeois and Linden Hu (Tufts Lyme Disease Initiative, Tufts University, Boston, MA) for generously providing the B31-AD strain propagated in *P. leucopus*. The contents are solely the responsibility of the authors and do not necessarily represent the official views of the National Institutes of Health.

## References

1. Mead P, Hinckley A, Kugeler K. 2024. Lyme Disease Surveillance and Epidemiology in the United States: A Historical Perspective. J Infect Dis 230:S11–S17.

2. Kugeler KJ, Earley A, Mead PS, Hinckley AF. 2024. Surveillance for Lyme Disease After Implementation of a Revised Case Definition - United States, 2022. MMWR Morb Mortal Wkly Rep 73:118–123.

3. Radolf JD, Caimano MJ, Stevenson B, Hu LT. 2012. Of ticks, mice and men: understanding the dual-host lifestyle of Lyme disease spirochaetes. Nat Rev Microbiol 10:87–99.

4. Radolf JD, Strle K, Lemieux JE, Strle F. 2021. Lyme Disease in Humans. Curr Issues Mol Biol 42:333–384.

5. Bobe JR, Jutras BL, Horn EJ, Embers ME, Bailey A, Moritz RL, Zhang Y, Soloski MJ, Ostfeld RS, Marconi RT, Aucott J, Ma’ayan A, Keesing F, Lewis K, Ben Mamoun C, Rebman AW, McClune ME, Breitschwerdt EB, Reddy PJ, Maggi R, Yang F, Nemser B, Ozcan A, Garner O, Di Carlo D, Ballard Z, Joung HA, Garcia-Romeu A, Griffiths RR, Baumgarth N, Fallon BA. 2021. Recent Progress in Lyme Disease and Remaining Challenges. Front Med (Lausanne) 8:666554.

6. Strle F, Strle K, Marques A, Henningsson AJ, Eikeland R, Lemieux JE, Tsao JI, Mead PS, Wormser GP. 2026. Lyme borreliosis. Nat Rev Dis Primers 12.

7. Baarsma ME, Hovius JW. 2024. Persistent Symptoms After Lyme Disease: Clinical Characteristics, Predictors, and Classification. J Infect Dis 230:S62–S69.

8. Wester KE, Nwokeabia BC, Hassan R, Dunphy T, Osondu M, Wonders C, Khaja M. 2024. What Makes It Tick: Exploring the Mechanisms of Post-treatment Lyme Disease Syndrome. Cureus 16:e64987.

9. Parthasarathy G, Gadila SKG. 2022. Neuropathogenicity of non-viable Borrelia burgdorferi ex vivo. Scientific Reports 12:688.

10. McClune ME, Ebohon O, Dressler JM, Davis MM, Tupik JD, Lochhead RB, Booth CJ, Steere AC, Jutras BL. 2025. The peptidoglycan of Borrelia burgdorferi can persist in discrete tissues and cause systemic responses consistent with chronic illness. Sci Transl Med 17:eadr2955.

11. Lantos PM, Wormser GP. 2014. Chronic Coinfections in Patients Diagnosed with Chronic Lyme Disease: A Systematic Review. The American Journal of Medicine 127:1105–1110.

12. Hajdusek O, Brangulis K, Robbertse L, Ghosh R, Carlo DD, Perner J. 2025. Lyme disease control 2.0: Advances and opportunities coming with Lyme disease vaccine VLA15. PLoS Pathog 21:e1013747.

13. Vogt NA, Sargeant JM, MacKinnon MC, Versluis AM. 2019. Efficacy of Borrelia burgdorferi vaccine in dogs in North America: A systematic review and meta-analysis. J Vet Intern Med 33:23–36.

14. O’Bier NS, Hatke AL, Camire AC, Marconi RT. 2021. Human and Veterinary Vaccines for Lyme Disease. Curr Issues Mol Biol 42:191–222.

15. Gomes-Solecki MJ, Brisson DR, Dattwyler RJ. 2006. Oral vaccine that breaks the transmission cycle of the Lyme disease spirochete can be delivered via bait. Vaccine 24:4440–9.

16. Gomes-Solecki M, Arnaboldi PM, Backenson PB, Benach JL, Cooper CL, Dattwyler RJ, Diuk-Wasser M, Fikrig E, Hovius JW, Laegreid W, Lundberg U, Marconi RT, Marques AR, Molloy P, Narasimhan S, Pal U, Pedra JHF, Plotkin S, Rock DL, Rosa P, Telford SR, Tsao J, Yang XF, Schutzer SE. 2020. Protective Immunity and New Vaccines for Lyme Disease. Clin Infect Dis 70:1768–1773.

17. Vannier E, Richer LM, Dinh DM, Brisson D, Ostfeld RS, Gomes-Solecki M. 2023. Deployment of a Reservoir-Targeted Vaccine Against Borrelia burgdorferi Reduces the Prevalence of Babesia microti Coinfection in Ixodes scapularis Ticks. J Infect Dis 227:1127–1131.

18. Bhattacharya D, Bensaci M, Luker KE, Luker G, Wisdom S, Telford SR, Hu LT. 2011. Development of a baited oral vaccine for use in reservoir-targeted strategies against Lyme disease. Vaccine 29:7818–25.

19. Scheckelhoff MR, Telford SR, Hu LT. 2006. Protective efficacy of an oral vaccine to reduce carriage of Borrelia burgdorferi (strain N40) in mouse and tick reservoirs. Vaccine 24:1949–57.

20. Rollend L, Fish D, Childs JE. 2013. Transovarial transmission of Borrelia spirochetes by Ixodes scapularis: a summary of the literature and recent observations. Ticks Tick Borne Dis 4:46–51.

21. Couret J, Dyer MC, Mather TN, Han S, Tsao JI, Lebrun RA, Ginsberg HS. 2017. Acquisition of Borrelia burgdorferi Infection by Larval Ixodes scapularis (Acari: Ixodidae) Associated With Engorgement Measures. J Med Entomol 54:1055–1060.

22. Meirelles Richer L, Aroso M, Contente-Cuomo T, Ivanova L, Gomes-Solecki M. 2011. Reservoir targeted vaccine for lyme borreliosis induces a yearlong, neutralizing antibody response to OspA in white-footed mice. Clin Vaccine Immunol 18:1809–16.

23. Balderrama-Gutierrez G, Milovic A, Cook VJ, Islam MN, Zhang Y, Kiaris H, Belisle JT, Mortazavi A, Barbour AG. 2021. An Infection-Tolerant Mammalian Reservoir for Several Zoonotic Agents Broadly Counters the Inflammatory Effects of Endotoxin. mBio 12.

24. Milovic A, Duong JV, Barbour AG. 2023. The white-footed deermouse, an infection-tolerant reservoir for several zoonotic agents, tempers interferon responses to endotoxin in comparison to the mouse and rat doi:10.7554/elife.90135.2. eLife Sciences Publications, Ltd.

25. Obaid MK, Islam N, Alouffi A, Khan AZ, da Silva Vaz I, Jr., Tanaka T, Ali A. 2022. Acaricides Resistance in Ticks: Selection, Diagnosis, Mechanisms, and Mitigation. Front Cell Infect Microbiol 12:941831.

26. Savage JDT, Moore CM. 2024. How do host population dynamics impact Lyme disease risk dynamics in theoretical models? PLoS One 19:e0302874.

27. Richer LM, Brisson D, Melo R, Ostfeld RS, Zeidner N, Gomes-Solecki M. 2014. Reservoir targeted vaccine against Borrelia burgdorferi: a new strategy to prevent Lyme disease transmission. J Infect Dis 209:1972–80.

28. Yamada M, Gomez JC, Chugh PE, Lowell CA, Dinauer MC, Dittmer DP, Doerschuk CM. 2011. Interferon-γ production by neutrophils during bacterial pneumonia in mice. Am J Respir Crit Care Med 183:1391–401.

29. Schwan TG, Piesman J, Golde WT, Dolan MC, Rosa PA. 1995. Induction of an outer surface protein on Borrelia burgdorferi during tick feeding. Proc Natl Acad Sci U S A 92:2909–13.

30. Steere AC, Sikand VK, Meurice F, Parenti DL, Fikrig E, Schoen RT, Nowakowski J, Schmid CH, Laukamp S, Buscarino C, Krause DS. 1998. Vaccination against Lyme disease with recombinant Borrelia burgdorferi outer-surface lipoprotein A with adjuvant. Lyme Disease Vaccine Study Group. N Engl J Med 339:209–15.

31. de Silva AM, Telford SR, 3rd, Brunet LR, Barthold SW, Fikrig E. 1996. Borrelia burgdorferi OspA is an arthropod-specific transmission-blocking Lyme disease vaccine. J Exp Med 183:271–5.

32. Bhatia B, Hillman C, Carracoi V, Cheff BN, Tilly K, Rosa PA. 2018. Infection history of the blood-meal host dictates pathogenic potential of the Lyme disease spirochete within the feeding tick vector. PLoS Pathog 14:e1006959.

33. Bockenstedt LK, Hodzic E, Feng S, Bourrel KW, de Silva A, Montgomery RR, Fikrig E, Radolf JD, Barthold SW. 1997. Borrelia burgdorferi strain-specific Osp C-mediated immunity in mice. Infect Immun 65:4661–7.

34. Earnhart CG, Buckles EL, Marconi RT. 2007. Development of an OspC-based tetravalent, recombinant, chimeric vaccinogen that elicits bactericidal antibody against diverse Lyme disease spirochete strains. Vaccine 25:466–80.

35. Earnhart CG, Marconi RT. 2007. An octavalent lyme disease vaccine induces antibodies that recognize all incorporated OspC type-specific sequences. Hum Vaccin 3:281–9.

36. Knobel DL, du Toit JT, Bingham J. 2002. Development of a bait and baiting system for delivery of oral rabies vaccine to free-ranging African wild dogs (Lycaon pictus). J Wildl Dis 38:352–62.

37. Brochier B, Kieny MP, Costy F, Coppens P, Bauduin B, Lecocq JP, Languet B, Chappuis G, Desmettre P, Afiademanyo K, et al. 1991. Large-scale eradication of rabies using recombinant vaccinia-rabies vaccine. Nature 354:520–2.

38. Slate D, Algeo TP, Nelson KM, Chipman RB, Donovan D, Blanton JD, Niezgoda M, Rupprecht CE. 2009. Oral rabies vaccination in north america: opportunities, complexities, and challenges. PLoS Negl Trop Dis 3:e549.

39. Rossi S, Staubach C, Blome S, Guberti V, Thulke HH, Vos A, Koenen F, Le Potier MF. 2015. Controlling of CSFV in European wild boar using oral vaccination: a review. Front Microbiol 6:1141.

40. Gomes-Solecki M, Richer L. 2018. Recombinant E. coli Dualistic Role as an Antigen-adjuvant Delivery Vehicle for Oral Immunization. Methods Mol Biol 1690:347–357.

41. Bensaci M, Bhattacharya D, Clark R, Hu LT. 2012. Oral vaccination with vaccinia virus expressing the tick antigen subolesin inhibits tick feeding and transmission of Borrelia burgdorferi. Vaccine 30:6040–6.

42. Rios S, Bhattachan B, Wirblich C, Chandwani A, Vavilikolanu K, Myers JF, Kitsou C, Pal U, Schnell MJ. 2025. Rabies virus-vectored Lyme disease vaccine provides long-term protection against tick-transmitted Borrelia burgdorferi. NPJ Vaccines 10:231.

43. Rios S, Bhattachan B, Vavilikolanu K, Kitsou C, Pal U, Schnell MJ. 2024. The Development of a Rabies Virus-Vectored Vaccine against Borrelia burgdorferi, Targeting BBI39. Vaccines (Basel) 12.

44. Stafford KC, 3rd, Williams SC, van Oosterwijk JG, Linske MA, Zatechka S, Richer LM, Molaei G, Przybyszewski C, Wikel SK. 2020. Field evaluation of a novel oral reservoir-targeted vaccine against Borrelia burgdorferi utilizing an inactivated whole-cell bacterial antigen expression vehicle. Exp Appl Acarol 80:257–268.

45. Kumaresan V, Smith TC, 2nd, Lumbreras M, Ingle TM, Kilgore N, Starling-Lin JF, Horn EJ, Seshu J. 2025. Protective efficacy of mutant strains of Borrelia burgdorferi as potential reservoir host-targeted biologics against Lyme disease. bioRxiv doi:10.1101/2025.07.09.663885.

46. Erdile LF, Brandt MA, Warakomski DJ, Westrack GJ, Sadziene A, Barbour AG, Mays JP. 1993. Role of attached lipid in immunogenicity of Borrelia burgdorferi OspA. Infect Immun 61:81–90.

47. Young TJ, Johnston KP, Mishima K, Tanaka H. 1999. Encapsulation of lysozyme in a biodegradable polymer by precipitation with a vapor-over-liquid antisolvent. J Pharm Sci 88:640–50.

48. Yang Q, Owusu-Ababio G. 2000. Biodegradable progesterone microsphere delivery system for osteoporosis therapy. Drug Dev Ind Pharm 26:61–70.

49. Pinero J, Temporal RM, Silva-Goncalves AJ, Jimenez IA, Bazzocchi IL, Oliva A, Perera A, Leon LL, Valladares B. 2006. New administration model of trans-chalcone biodegradable polymers for the treatment of experimental leishmaniasis. Acta Trop 98:59–65.

50. Ahmed KK, Geary SM, Salem AK. 2016. Development and Evaluation of Biodegradable Particles Coloaded With Antigen and the Toll-Like Receptor Agonist, Pentaerythritol Lipid A, as a Cancer Vaccine. J Pharm Sci 105:1173–9.

51. Ahmed TA, Aljaeid BM. 2016. Preparation, characterization, and potential application of chitosan, chitosan derivatives, and chitosan metal nanoparticles in pharmaceutical drug delivery. Drug Des Devel Ther 10:483–507.

52. Gala RP, D’Souza M, Zughaier SM. 2016. Evaluation of various adjuvant nanoparticulate formulations for meningococcal capsular polysaccharide-based vaccine. Vaccine 34:3260–7.

53. Hiremath J, Kang KI, Xia M, Elaish M, Binjawadagi B, Ouyang K, Dhakal S, Arcos J, Torrelles JB, Jiang X, Lee CW, Renukaradhya GJ. 2016. Entrapment of H1N1 Influenza Virus Derived Conserved Peptides in PLGA Nanoparticles Enhances T Cell Response and Vaccine Efficacy in Pigs. PLoS One 11:e0151922.

54. Kokate RA, Chaudhary P, Sun X, Thamake SI, Maji S, Chib R, Vishwanatha JK, Jones HP. 2016. Rationalizing the use of functionalized poly-lactic-co-glycolic acid nanoparticles for dendritic cell-based targeted anticancer therapy. Nanomedicine (Lond) 11:479–94.

55. Lee BK, Yun Y, Park K. 2016. PLA micro- and nano-particles. Adv Drug Deliv Rev 107:176–191.

56. Silva AL, Soema PC, Slutter B, Ossendorp F, Jiskoot W. 2016. PLGA particulate delivery systems for subunit vaccines: Linking particle properties to immunogenicity. Hum Vaccin Immunother 12:1056–69.

57. Varypataki EM, Silva AL, Barnier-Quer C, Collin N, Ossendorp F, Jiskoot W. 2016. Synthetic long peptide-based vaccine formulations for induction of cell mediated immunity: A comparative study of cationic liposomes and PLGA nanoparticles. J Control Release 226:98–106.

58. Zhang NZ, Xu Y, Wang M, Chen J, Huang SY, Gao Q, Zhu XQ. 2016. Vaccination with Toxoplasma gondii calcium-dependent protein kinase 6 and rhoptry protein 18 encapsulated in poly(lactide-co-glycolide) microspheres induces long-term protective immunity in mice. BMC Infect Dis 16:168.

59. McGhee JR, Fujihashi K, Xu-Amano J, Jackson RJ, Elson CO, Beagley KW, Kiyono H. 1993. New perspectives in mucosal immunity with emphasis on vaccine development. Semin Hematol 30:3–12; discussion 13-5.

60. Jackson RJ, Fujihashi K, Xu-Amano J, Kiyono H, Elson CO, McGhee JR. 1993. Optimizing oral vaccines: induction of systemic and mucosal B-cell and antibody responses to tetanus toxoid by use of cholera toxin as an adjuvant. Infect Immun 61:4272–9.

61. Fujihashi K, Kato H, van Ginkel FW, Koga T, Boyaka PN, Jackson RJ, Kato R, Hagiwara Y, Etani Y, Goma I, Fujihashi K, Kiyono H, McGhee JR. 2001. A revisit of mucosal IgA immunity and oral tolerance. Acta Odontol Scand 59:301–8.

62. van Ginkel FW, Jackson RJ, Yuki Y, McGhee JR. 2000. Cutting edge: the mucosal adjuvant cholera toxin redirects vaccine proteins into olfactory tissues. J Immunol 165:4778–82.

63. Nakamura Y, Kimura S, Hase K. 2018. M cell-dependent antigen uptake on follicle-associated epithelium for mucosal immune surveillance. Inflamm Regen 38:15.

64. Hori M, Onishi H, Machida Y. 2005. Evaluation of Eudragit-coated chitosan microparticles as an oral immune delivery system. Int J Pharm 297:223–34.

65. Uddin MJ, Gill HS. 2017. Ragweed pollen as an oral vaccine delivery system: Mechanistic insights. J Control Release 268:416–426.

66. Tsao JI, Hamer SA, Han S, Sidge JL, Hickling GJ. 2021. The Contribution of Wildlife Hosts to the Rise of Ticks and Tick-Borne Diseases in North America. J Med Entomol 58:1565–1587.

67. Bunikis J, Tsao J, Luke CJ, Luna MG, Fish D, Barbour AG. 2004. Borrelia burgdorferi infection in a natural population of Peromyscus Leucopus mice: a longitudinal study in an area where Lyme Borreliosis is highly endemic. J Infect Dis 189:1515–23.

68. Petzke MM, Iyer R, Love AC, Spieler Z, Brooks A, Schwartz I. 2016. Borrelia burgdorferi induces a type I interferon response during early stages of disseminated infection in mice. BMC Microbiol 16:29.

69. Kim S-H, Seo K-W, Kim J, Lee K-Y, Jang Y-S. 2010. The M Cell-Targeting Ligand Promotes Antigen Delivery and Induces Antigen-Specific Immune Responses in Mucosal Vaccination. The Journal of Immunology 185:5787–5795.

70. Wells Christopher C, Petnicki-Ocwieja T, Tan S, Bunnell Stephen C, Telford Sam R, Hu Linden T, Bourgeois Jeffrey S. 2025. Differentiating Peromyscus leucopus bone marrow-derived macrophages for characterization of responses to Borrelia burgdorferi and lipopolysaccharide. Infection and Immunity 93:e00581–24.

71. Eisen RJ, Eisen L, Beard CB. 2016. County-Scale Distribution of Ixodes scapularis and Ixodes pacificus (Acari: Ixodidae) in the Continental United States. J Med Entomol 53:349–86.

72. Ostfeld RS, Canham CD, Oggenfuss K, Winchcombe RJ, Keesing F. 2006. Climate, deer, rodents, and acorns as determinants of variation in lyme-disease risk. PLoS Biol 4:e145.

73. Kumaresan V, Pahari S, Hung C-Y, Hermann BP, Schlesinger LS, Seshu J. 2026. Role of dual specificity phosphatase 1 in influencing inflammatory pathways in macrophages modulated by Borrelia burgdorferi lipoproteins. Frontiers in Immunology Volume 17–2026.

74. Bernard Q, Phelan JP, Hu LT. 2020. Controlling Lyme Disease: New Paradigms for Targeting the Tick-Pathogen-Reservoir Axis on the Horizon. Front Cell Infect Microbiol 10:607170.

75. Ou B, Yang Y, Lv H, Lin X, Zhang M. 2023. Current Progress and Challenges in the Study of Adjuvants for Oral Vaccines. BioDrugs 37:143–180.

76. Holmgren J, Lycke N, Czerkinsky C. 1993. Cholera toxin and cholera B subunit as oral-mucosal adjuvant and antigen vector systems. Vaccine 11:1179–84.

77. Baldauf KJ, Royal JM, Kouokam JC, Haribabu B, Jala VR, Yaddanapudi K, Hamorsky KT, Dryden GW, Matoba N. 2017. Oral administration of a recombinant cholera toxin B subunit promotes mucosal healing in the colon. Mucosal Immunology 10:887–900.

78. Gloudemans AK, Plantinga M, Guilliams M, Willart MA, Ozir-Fazalalikhan A, van der Ham A, Boon L, Harris NL, Hammad H, Hoogsteden HC, Yazdanbakhsh M, Hendriks RW, Lambrecht BN, Smits HH. 2013. The Mucosal Adjuvant Cholera Toxin B Instructs Non-Mucosal Dendritic Cells to Promote IgA Production Via Retinoic Acid and TGF-β. PLOS ONE 8:e59822.

79. McCarty M, Hernandez SA, Malfetano J, Leao AC, Villar MJ, Yang X, Chen YL, Lee J, Liu Z, Strych U, Bottazzi ME, Pal U, Strle K, Chen WH, Lin YP. 2026. Develop a durable, memory-driven, CspZ-targeting Lyme disease vaccine by rationale adjuvant selection. bioRxiv doi:10.64898/2026.01.08.698480.

80. Marcinkiewicz AL, Lieknina I, Yang X, Lederman PL, Hart TM, Yates J, Chen WH, Bottazzi ME, Mantis NJ, Kraiczy P, Pal U, Tars K, Lin YP. 2020. The Factor H-Binding Site of CspZ as a Protective Target against Multistrain, Tick-Transmitted Lyme Disease. Infect Immun 88.

81. Xiao S, Kumar M, Yang X, Akkoyunlu M, Collins PL, Samal SK, Pal U. 2011. A host-restricted viral vector for antigen-specific immunization against Lyme disease pathogen. Vaccine 29:5294–303.

82. Brangulis K, Malfetano J, Marcinkiewicz AL, Wang A, Chen YL, Lee J, Liu Z, Yang X, Strych U, Tupina D, Akopjana I, Bottazzi ME, Pal U, Hsieh CL, Chen WH, Lin YP. 2025. Mechanistic insights into the structure-based design of a CspZ-targeting Lyme disease vaccine. Nat Commun 16:2898.

83. Chen YL, Lee J, Liu Z, Strych U, Bottazzi ME, Lin YP, Chen WH. 2024. Biophysical and biochemical characterization of a recombinant Lyme disease vaccine antigen, CspZ-YA. Int J Biol Macromol 259:129295.

84. Brangulis K, Malfetano J, Marcinkiewicz AL, Wang A, Chen YL, Lee J, Liu Z, Yang X, Strych U, Bottazzi ME, Pal U, Hsieh CL, Chen WH, Lin YP. 2024. Mechanistic insights into structure-based design of a Lyme disease subunit vaccine. bioRxiv doi:10.1101/2024.10.23.619738.

85. Chen YL, Marcinkiewicz AL, Nowak TA, Tyagi Kundu R, Liu Z, Strych U, Bottazzi ME, Chen WH, Lin YP. 2022. CspZ FH-Binding Sites as Epitopes Promote Antibody-Mediated Lyme Borreliae Clearance. Infect Immun 90:e0006222.

86. Chen WH, Strych U, Bottazzi ME, Lin YP. 2022. Past, present, and future of Lyme disease vaccines: antigen engineering approaches and mechanistic insights. Expert Rev Vaccines 21:1405–1417.

87. Melo R, Richer L, Johnson DL, Gomes-Solecki M. 2016. Oral Immunization with OspC Does Not Prevent Tick-Borne Borrelia burgdorferi Infection. PLoS One 11:e0151850.

88. Gomes-Solecki M. 2014. Blocking pathogen transmission at the source: reservoir targeted OspA-based vaccines against Borrelia burgdorferi. Front Cell Infect Microbiol 4:136.

89. Hou J, Liu Y, Hsi J, Wang H, Tao R, Shao Y. 2014. Cholera toxin B subunit acts as a potent systemic adjuvant for HIV-1 DNA vaccination intramuscularly in mice. Hum Vaccin Immunother 10:1274–83.

90. Tsao JI, Wootton JT, Bunikis J, Luna MG, Fish D, Barbour AG. 2004. An ecological approach to preventing human infection: vaccinating wild mouse reservoirs intervenes in the Lyme disease cycle. Proc Natl Acad Sci U S A 101:18159–64.

91. Lundberg U, Hochreiter R, Timofoyeva Y, Kanevsky I, Meinke A, Anderson AS, Simon R. 2024. Preclinical Evidence for the Protective Capacity of Antibodies Induced by Lyme Vaccine Candidate VLA15 in People. Open Forum Infect Dis 11:ofae467.

92. Barbour AG. 2017. Infection resistance and tolerance in Peromyscus spp., natural reservoirs of microbes that are virulent for humans. Semin Cell Dev Biol 61:115–122.

93. Tsao J, Barbour AG, Luke CJ, Fikrig E, Fish D. 2001. OspA immunization decreases transmission of Borrelia burgdorferi spirochetes from infected Peromyscus leucopus mice to larval Ixodes scapularis ticks. Vector Borne Zoonotic Dis 1:65–74.

94. Rogovskyy AS, Pliasas VC, Buhrer R, Lewy K, Wiener DJ, Jung Y, Bova J, Rogovska Y, Kim SJ, Jeon E. 2024. Do white-footed mice, the main reservoir of the Lyme disease pathogen in the United States, clinically respond to the borrelial tenancy? Infect Immun 92:e0038224.

95. Gaber AM, Mandric I, Nitirahardjo C, Piontkivska H, Hillhouse AE, Threadgill DW, Zelikovsky A, Rogovskyy AS. 2023. Comparative transcriptome analysis of Peromyscus leucopus and C3H mice infected with the Lyme disease pathogen. Front Cell Infect Microbiol 13:1115350.

96. Rogovskyy AS, Casselli T, Tourand Y, Jones CR, Owen JP, Mason KL, Scoles GA, Bankhead T. 2015. Evaluation of the Importance of VlsE Antigenic Variation for the Enzootic Cycle of Borrelia burgdorferi. PLoS One 10:e0124268.

97. Narasimhan S, Fish D, Pedra JHF, Pal U, Fikrig E. 2023. A ticking time bomb hidden in plain sight. Sci Transl Med 15:eadi7829.

98. Joo SY, Lee SJ, Lee SJ, Seo SA, Kim C-W, Hong S-H, Kim I, Kim M. 2025. Single-cell RNA sequencing reveals that a high salt diet increases IL6R-JAK1-STAT3 gene signaling pathway activity in intestinal B cells and Tfh cells of salt-sensitive hypertension animal model. Scientific Reports 15:25756.

99. Dar HY, Singh A, Shukla P, Anupam R, Mondal RK, Mishra PK, Srivastava RK. 2018. High dietary salt intake correlates with modulated Th17-Treg cell balance resulting in enhanced bone loss and impaired bone-microarchitecture in male mice. Scientific Reports 8:2503.

100. Luo M, Shao B, Yu JY, Liu T, Liang X, Lu L, Ye TH, He ZY, Xiao HY, Wei XW. 2017. Simultaneous enhancement of cellular and humoral immunity by the high salt formulation of Al(OH)(3) adjuvant. Cell Res 27:586–589.

101. Johnson EE, Hart TM, Fikrig E. 2024. Vaccination to Prevent Lyme Disease: A Movement Towards Anti-Tick Approaches. The Journal of Infectious Diseases 230:S82–S86.

102. Elias AF, Stewart PE, Grimm D, Caimano MJ, Eggers CH, Tilly K, Bono JL, Akins DR, Radolf JD, Schwan TG, Rosa P. 2002. Clonal polymorphism of Borrelia burgdorferi strain B31 MI: implications for mutagenesis in an infectious strain background. Infect Immun 70:2139–50.

103. Bourgeois JS, McCarthy JE, Turk S-P, You SS, Bernard Q, Clendenen LH, Wormser GP, Marcos LA, Dardick K, Telford SR, Marques AR, Hu LT. 2025. *Peromyscus leucopus*, *Mus musculus*, and humans have distinct transcriptomic responses to larval *Ixodes scapularis* bites. Infection and Immunity 93:e00065–25.

104. Raju BV, Esteve-Gassent MD, Karna SL, Miller CL, Van Laar TA, Seshu J. 2011. Oligopeptide permease A5 modulates vertebrate host-specific adaptation of Borrelia burgdorferi. Infect Immun 79:3407–20.

105. Van Laar TA, Lin YH, Miller CL, Karna SL, Chambers JP, Seshu J. 2012. Effect of levels of acetate on the mevalonate pathway of Borrelia burgdorferi. PLoS One 7:e38171.

106. Van Laar TA, Hole C, Rajasekhar Karna SL, Miller CL, Reddick R, Wormley FL, Seshu J. 2016. Statins reduce spirochetal burden and modulate immune responses in the C3H/HeN mouse model of Lyme disease. Microbes Infect 18:430–435.

107. Karna SL, Sanjuan E, Esteve-Gassent MD, Miller CL, Maruskova M, Seshu J. 2011. CsrA modulates levels of lipoproteins and key regulators of gene expression critical for pathogenic mechanisms of Borrelia burgdorferi. Infect Immun 79:732–44.

108. Karna SL, Prabhu RG, Lin YH, Miller CL, Seshu J. 2013. Contributions of environmental signals and conserved residues to the functions of carbon storage regulator A of Borrelia burgdorferi. Infect Immun 81:2972–85.

109. Miller CL, Karna SL, Seshu J. 2013. Borrelia host adaptation Regulator (BadR) regulates rpoS to modulate host adaptation and virulence factors in Borrelia burgdorferi. Mol Microbiol 88:105–24.

110. Caputo A, Castaldello A, Brocca-Cofano E, Voltan R, Bortolazzi F, Altavilla G, Sparnacci K, Laus M, Tondelli L, Gavioli R, Ensoli B. 2009. Induction of humoral and enhanced cellular immune responses by novel core-shell nanosphere- and microsphere-based vaccine formulations following systemic and mucosal administration. Vaccine 27:3605–15.

111. Chen Y, Vargas SM, Smith TC, 2nd, Karna SLR, MacMackin Ingle T, Wozniak KL, Wormley FL, Jr., Seshu J. 2021. Borrelia peptidoglycan interacting Protein (BpiP) contributes to the fitness of Borrelia burgdorferi against host-derived factors and influences virulence in mouse models of Lyme disease. PLoS Pathog 17:e1009535.

112. Bourgeois JS, You SS, Clendenen LH, Shrestha M, Petnicki-Ocwieja T, Telford SR, 3rd, Hu LT. 2024. Comparative reservoir competence of Peromyscus leucopus, C57BL/6J, and C3H/HeN for Borrelia burgdorferi B31. Appl Environ Microbiol 90:e0082224.

